# Rapid DNA unwinding accelerates genome editing by engineered CRISPR-Cas9

**DOI:** 10.1101/2023.12.14.571777

**Authors:** Amy R. Eggers, Kai Chen, Katarzyna M. Soczek, Owen T. Tuck, Erin E. Doherty, Brittney W. Thornton, Bryant Xu, Marena I. Trinidad, Jennifer A. Doudna

**Affiliations:** Department of Molecular and Cell Biology, University of California, Berkeley; Berkeley, CA, USA; Innovative Genomics Institute; University of California, Berkeley, CA, USA; Howard Hughes Medical Institute, University of California, Berkeley; Berkeley CA, USA; Gladstone Institutes; San Francisco, CA, USA; Gladstone-UCSF Institute of Genomic Immunology; San Francisco, CA, USA; Molecular Biophysics and Integrated Bioimaging Division, Lawrence Berkeley National Laboratory; Berkeley, CA, USA; Department of Chemistry, University of California, Berkeley; Berkeley, CA, USA; California Institute for Quantitative Biosciences, University of California, Berkeley; Berkeley, CA, USA

## Abstract

Thermostable CRISPR-Cas9 enzymes could improve genome editing efficiency and delivery due to extended protein lifetimes. However, initial experimentation demonstrated Geobacillus stearothermophilus Cas9 (GeoCas9) to be virtually inactive when used in cultured human cells. Laboratory-evolved variants of GeoCas9 overcome this natural limitation by acquiring mutations in the wedge (WED) domain that produce >100-fold higher genome editing levels. Cryo-EM structures of the wildtype and improved GeoCas9 (iGeoCas9) enzymes reveal extended contacts between the WED domain of iGeoCas9 and DNA substrates. Biochemical analysis shows that iGeoCas9 accelerates DNA unwinding to capture substrates under the magnesium-restricted conditions typical of mammalian but not bacterial cells. These findings enabled rational engineering of other Cas9 orthologs to enhance genome editing levels, pointing to a general strategy for editing enzyme improvement. Together, these results uncover a new role for the Cas9 WED domain in DNA unwinding and demonstrate how accelerated target unwinding dramatically improves Cas9-induced genome editing activity.

## INTRODUCTION

Clustered regularly interspaced short palindromic repeats (CRISPR) and CRISPR-associated (Cas) proteins provide adaptive immunity for prokaryotes by selectively targeting and cleaving nucleic acids of invading viruses and mobile genetic elements^1^. This defense mechanism relies on CRISPR RNA-guided Cas proteins to distinguish foreign DNA from endogenous DNA. The ∼20-nucleotide guide sequence in the RNA uses base-pairing complementarity to recognize a foreign DNA target, triggering its Cas-mediated cleavage^2^. The ease of guide RNA reprogramming is central to applications of CRISPR-Cas systems for genome editing^3,4^. CRISPR-Cas9 from *Streptococcus pyogenes* (SpyCas9), the most widely adopted editor, is commonly used in research as well as clinical and agricultural applications^5^. However, its sensitivity to aggregation or proteolytic inactivation can inhibit genome editing outcomes under some conditions *in vitro* and *in vivo*^6^.

The thermostable enzyme GeoCas9^7,8^ was hypothesized to have enhanced genome editing activity based on extended protein lifetime, but this was not borne out by experimental evidence. Instead, data showed GeoCas9 to be a poor editor in human cells despite its thermostability and robust biochemical activity. Using directed evolution, GeoCas9 variants were identified that dramatically improve genome editing both in cultured cells and in mouse tissues^9^. The resulting iGeoCas9 enzymes, which show >100-fold increases in genome editing efficiencies, contain mutations in the wedge (WED) domain that increase activity while maintaining thermal tolerance and protein stability^9^.

To determine the molecular basis for the dramatic improvement of iGeoCas9 as a genome editor, we used a combination of cryo-electron microscopy (cryo-EM) and biochemical analysis to compare the structures and behaviors of wildtype versus iGeoCas9 enzymes. The three WED domain mutations responsible for improved editing by iGeoCas9 were found to establish new interactions with target double-stranded DNA (dsDNA), leading to enhanced DNA binding and a relaxed preference for protospacer-adjacent motif (PAM) base pairs next to the target sequence. Using a fluorescent reporter assay, we found that the improved dsDNA binding dramatically accelerated DNA unwinding, expanding the target sequence space by reducing PAM specificity. Furthermore, iGeoCas9 WED-domain mutations enable the enzyme to function at dramatically reduced magnesium ion concentrations consistent with those found in mammalian but not bacterial cells. Similar WED-domain mutations introduced into other Cas9 enzymes also enhanced genome editing activity in cells, implicating this structural region as a pivotal and previously unappreciated moderator of Cas9 target recognition. Together, these data reveal an unexpected connection between protein-DNA binding and helix unwinding that explains how Cas9 enzymatic activity can be modified to enhance genome editing.

## RESULTS

### GeoCas9 molecular structures reveal unexpected WED domain-DNA contacts

To understand why iGeoCas9 is an efficient genome editor when its parent enzyme is not, we first explored the structural differences between wildtype GeoCas9 and iGeoCas9. iGeoCas9 contains 8 engineered mutations to wildtype GeoCas9 (Fig. 1A), including three in the Rec domain (E149G, T182I, N206D), one in the RuvC domain (P466Q), one in the phosphate lock loop (Q817R), and three in the WED domain (E843K, E884G, and K908R). In order to see how the mutations affect Cas9’s function on target DNA, we reconstituted the ternary structures of catalytically-deactivated wildtype GeoCas9 and iGeoCas9 (with nuclease-deactivating mutations, D8A and H582A, introduced to the RuvC and HNH domains, respectively) complexed with single-guide RNA (sgRNA) and target DNA to capture nucleic acid interactions that occur upon R-loop formation (Fig. 1B). Cryo-EM reconstructions of wildtype GeoCas9 and iGeoCas9 (3.17 Å and 2.63 Å resolution, respectively), reveal similar domain architecture to other type II-C enzymes, with a smaller recognition (REC) domain relative to type II-A enzymes (Fig. 1C; Figs. S1, S2). A full 21-nt target strand (TS) DNA binds the complementary guide RNA sequence and the entire 5’-N_4_CAAA-3’ PAM-containing dsDNA duplex. Only one nucleotide of the non-target strand (NTS) was resolved, with the remainder disordered. The sgRNA scaffold forms a triple stem-loop architecture, with a nucleotide triplex in stem loop 3; unsharpened maps indicate a possible fourth stem loop, but the density was insufficient to build a reliable model (Fig. 1D). Wildtype GeoCas9 and iGeoCas9 structures are overall very similar, both having disordered HNH domains, likely representing the pre-catalytic states (Fig. S3).

**Figure 1.**
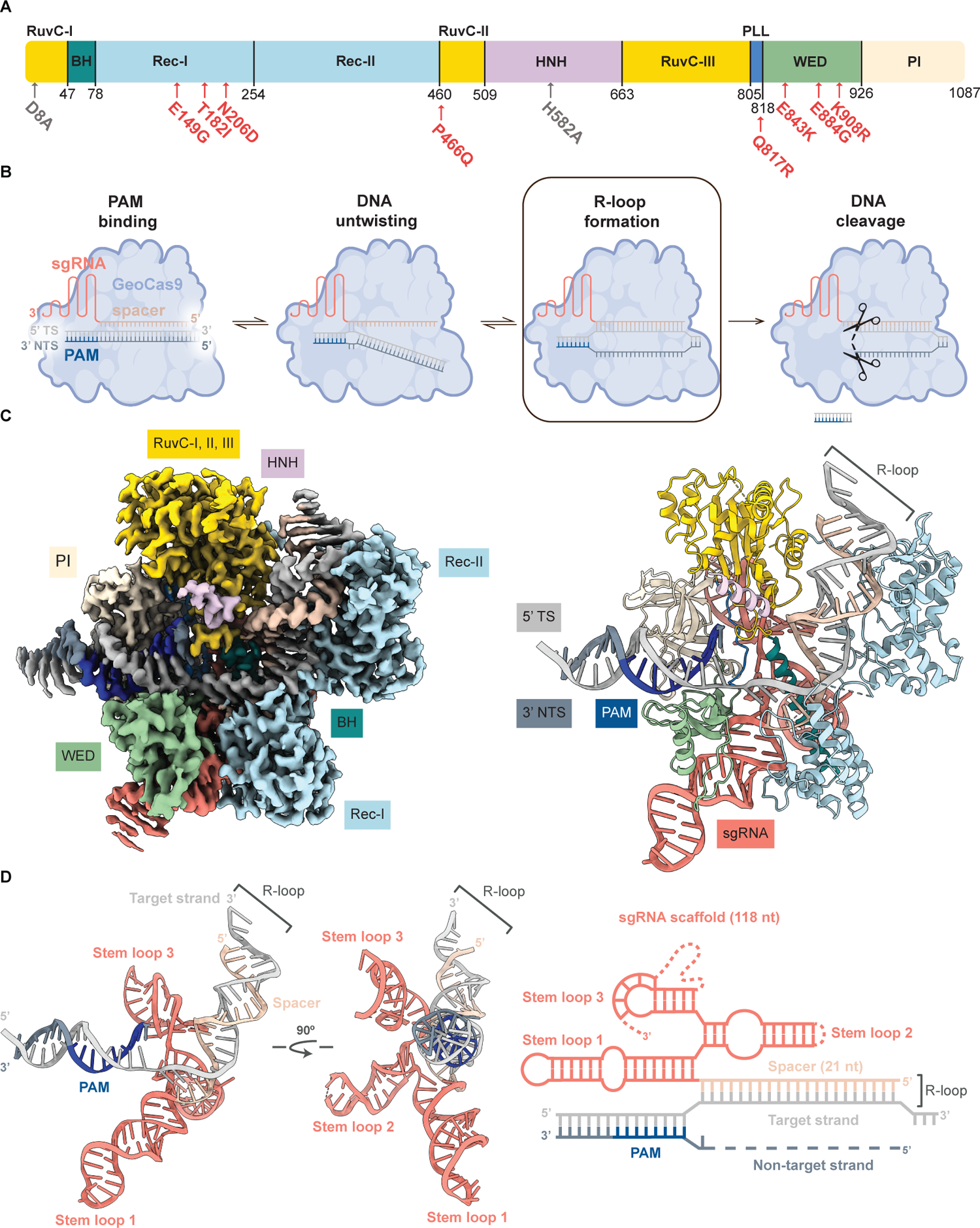
Cryo-EM iGeoCas9-sgRNA-DNA ternary complex. **(A)** Domain organization of GeoCas9. iGeoCas9 mutations indicated with red arrows. Deactivating mutations indicated with gray arrows. **(B)** CRISPR-Cas9 dsDNA targeting pathway for active enzymes. Deactivated Cas9 concludes at the R-loop formation step. **(C)** Surface (left) and ribbon (right) representation of iGeoCas9-sgRNA-DNA ternary complex. Domains are colored as in **1A**. Nucleic acid is colored as in **1D**. BH, bridge helix. WED, wedge domain. PI, PAM interacting domain. TS, target strand. NTS, non-target strand. PAM, protospacer adjacent motif. sgRNA, single guide RNA. See also Figure S1, S2, Table S1. **(D)** Cartoon representation of iGeoCas9 sgRNA-dsDNA complex (left) schematic representation of iGeoCas9 sgRNA:target DNA complex (right).

Prior studies demonstrated that the three WED domain mutations, E843K, E884G and K908R, in iGeoCas9 had the biggest impact on promoting genome editing activity, but the mechanism was unclear^9^. The ∼100 amino acid WED domain comprises four alpha helices and two beta sheets that recognize the repeat:anti-repeat region of the sgRNA and the dsDNA located upstream of the target region (Fig. 2A).

**Figure 2.**
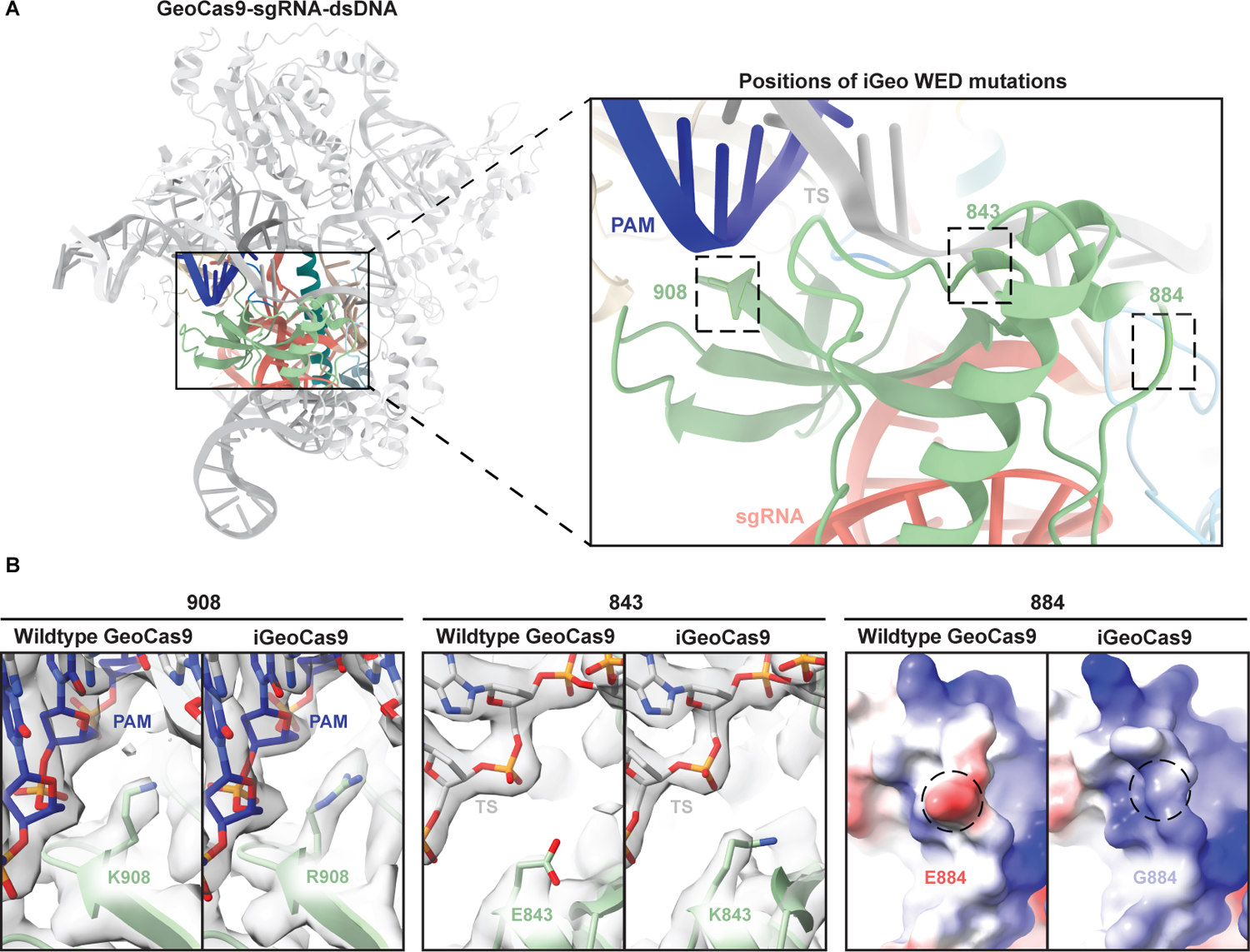
Structural comparison of wildtype GeoCas9 and iGeoCas9 WED domain. **(A)** Ribbon representation of iGeoCas9 WED domain. Amino acid mutation positions in iGeoCas9 are boxed with a dashed line. **(B)** Comparison of wildtype GeoCas9 and iGeoCas9 at WED domain amino acid positions 908, 843, and 884. Positions 908 and 843 are represented by an electron density map. Position 884 is represented by an electrostatic surface potential map (position 884 encircled with a dashed line).

The iGeoCas9 WED domain mutations (E843K, E884G and K908R) may alter contacts with the phosphate backbone of adjacent target-strand DNA backbone. Indeed, the positively charged lysine (K) at position 843 in iGeoCas9 is situated to establish a new electrostatic interaction with the negatively charged phosphate backbone of TS DNA (Fig. 2B). Mutation K908R in iGeoCas9, located in the minor groove adjacent to the NTS, potentially enhances DNA binding electrostatics (Fig. 2B). Together, mutations E843K and K908R in iGeoCas9 may augment DNA strand separation and R-loop formation required for DNA cleavage. While not in contact with nucleic acid in our experimental structure, the glutamate 884 to glycine mutation (E884G) in iGeoCas9 may further enhance nucleic acid interactions through altering the electrostatics (Fig. 2B). Based on these structural observations, we hypothesize the three WED domain mutations in iGeoCas9 favor DNA strand separation prior to cleavage. This motivated us to biochemically investigate how these mutations affect each individual step of the Cas9 catalytic pathway (Fig. 1B).

### iGeoCas9-catalyzed DNA cleavage has a relaxed PAM requirement

Cas9 interacts with an adjacent protospacer motif (PAM) before it can proceed with DNA target strand recognition and cleavage, the necessary precursor to genome editing^10^. To determine how the mutations in iGeoCas9 promote genome editing, we first investigated their effect on PAM sequence recognition. The PAM was previously determined as 5’-N_4_CRAA-3’ (where R = A/G, N = A/T/C/G) for wildtype GeoCas9^7^. Using a 6-carboxyfluorescein (6-FAM) labeled 60-base pair DNA substrate, we observed that wildtype GeoCas9 and iGeoCas9 have similar DNA cleavage activities against the native 5’-N_4_CAAA PAM (Fig. 3A and 3B). When dsDNA bearing different PAM sequences were employed as substrates, we found that iGeoCas9 has a much wider tolerance for non-native PAMs (5’-N_4_CAGA-3’, 5’-N_4_GCAA-3’ and 5’-N_4_TAAA-3’) compared to wildtype GeoCas9 (Fig. 3B and S4). These observations show that iGeoCas9 has acquired the ability to accommodate more diverse PAM sequences, allowing it to target a broader range of substrate sequences.

**Figure 3.**
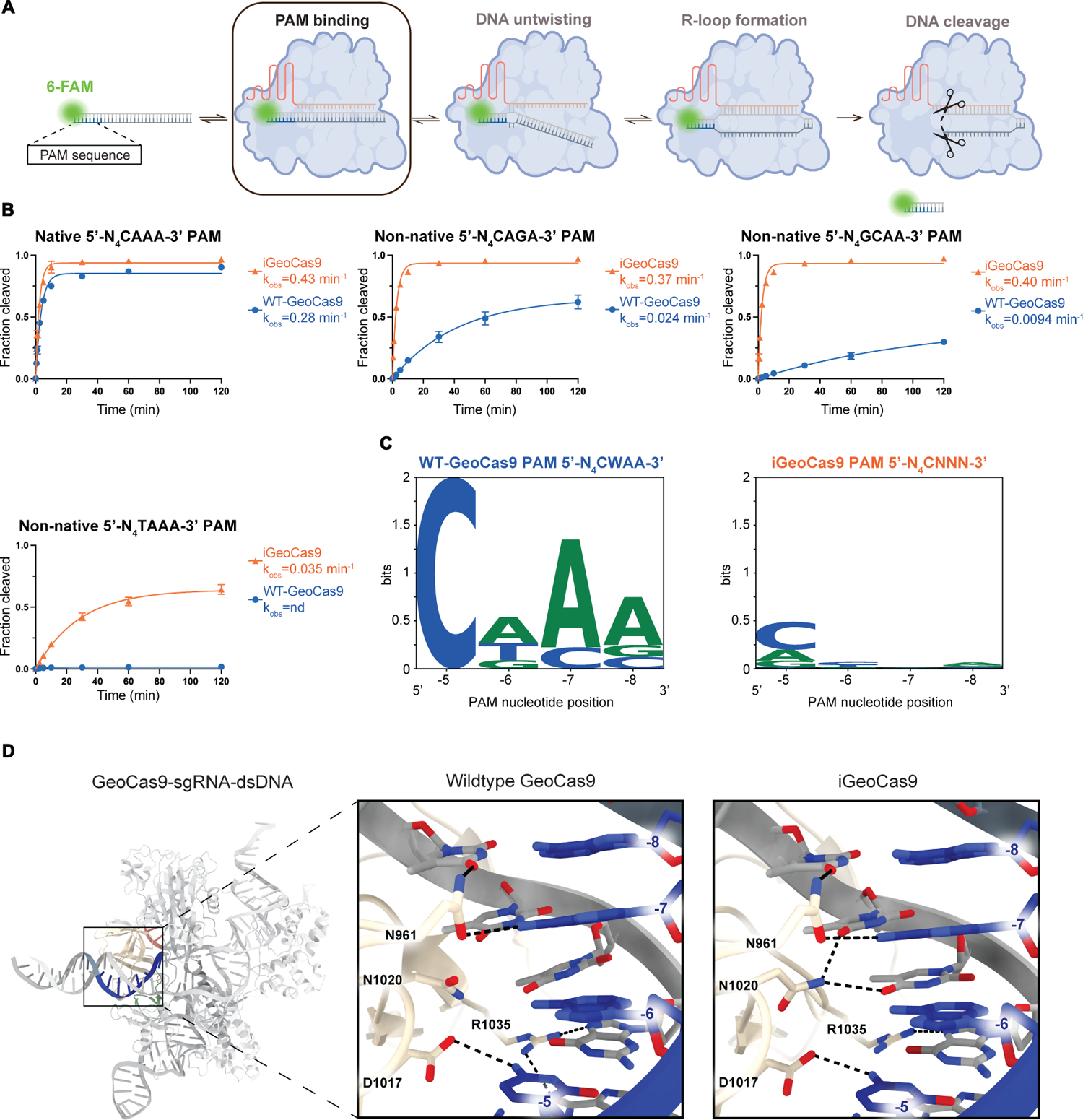
iGeoCas9 demonstrates enhanced activity targeting wildtype GeoCas9 non-native PAMs. **(A)** Cas9 cleavage reaction using 60 nt 5’ 6-FAM labeled dsDNA substrates with different PAM sequences. **(B)** *In vitro* dsDNA cleavage activity of wildtype (WT-)GeoCas9 and iGeoCas9 determined by denaturing PAGE (n = 3, mean ± SD). PAM contained in each substrate indicated above the graph. Fractions were collected at 0 sec, 30 sec, 1 min, 2.5 min, 5 min, 10 min, 30 min, 1 h, and 2 h. The k_obs_ for each Cas9 is listed in the sample legend. See also Figure S4. **(C)** WebLogo for sequences depleted from the PAM library. PAM position on X-axis and is numbered in the 5’ to 3’ direction in descending order from −1 to −8. G, C and T, blue; and A and G, green. **(D)** PAM nucleobase interacting amino acids (D961, N1020, D1017, and R1035) of wildtype GeoCas9 and iGeoCas9. PAM sequence position indicated adjacent to nucleotide. H-bond prediction was performed in ChimeraX with a distance tolerance of 0.400 Å and angle tolerance of 20 degrees. Alternate rotamer conformations were observed for N1020 and R1035 in wildtype GeoCas9 and iGeoCas9.

Next, we used a PAM depletion assay to compare the behavior of wildtype GeoCas9 and iGeoCas9 more comprehensively^11^. Purified guide RNA-Cas9 ribonucleoprotein (RNP) complexes were incubated with a plasmid library containing a PAM with four randomized nucleotides at positions −5 to −8 (5’-TTTTN_4_-3’). Successful PAM recognition depletes those sequences from the library relative to a non-targeting guide RNP control. Next-generation sequencing (NGS) revealed wildtype GeoCas9 was constricted to a consensus sequence of 5’-N_4_CWAA-3’ (where W= A/T), consistent with our prior report^7^. PAM depletion results showed that iGeoCas9 uses a relaxed PAM consensus sequence of 5’-N_4_CNNN-3’ (Fig. 3C). Although PAM specificity can be altered for different Cas proteins when mutations are introduced to the PAM-interacting (PI) domain^12,13^, iGeoCas9 does not contain mutations in the PI domain (Fig. 3D). PAM expansion is likely driven by the DNA backbone interactions in the WED domain ^14^. Together, these observations suggest that downstream steps in DNA unwinding or catalysis contribute to the expanded PAM compatibility of iGeoCas9.

### iGeoCas9 has improved DNA melting kinetics

The cryo-EM structures of wildtype GeoCas9 and iGeoCas9 suggest that mutations K908R and E843K in iGeoCas9 may enhance DNA backbone interactions on the non-target and target strands of the recognition sequence. To test whether the effect of mutations is to improve DNA unwinding upon Cas9-guide target binding, we used DNA substrates containing either perfectly matched (linear) or mismatched (mm) base pairs immediately adjacent to the PAM sequence (Fig. 4A). Previous studies demonstrated that type II-C Cas9s, have improved DNA unwinding and DNA cleavage kinetics with a thermodynamically destabilized DNA substrate containing two base pair mismatches adjacent to the PAM^15^. These mismatches assist in the initial disruption of the dsDNA helix, or DNA melting, that accompanies R-loop formation (Fig. 4A). To test whether the WED domain mutations change the ability of GeoCas9 to recognize and cleave DNA target sequences that have a non-native PAM, we designed a 60-bp DNA substrate bearing a PAM sequence, 5’-N_4_GAAA-3’, that is disfavored by wildtype GeoCas9. To further explore the effect of WED domain mutations, activities of wildtype GeoCas9 and iGeoCas9 were compared to an intermediate mutant from iGeoCas9’s laboratory evolutionary lineage, GeoCas9(R1), that does not contain the WED domain mutations (Fig. 4B). Under substrate-limiting (single-turnover) conditions with a linear substrate, initial rates for both the wildtype GeoCas9 and GeoCas9(R1) were similar (0.040 vs 0.055 min^-1^), suggesting the mutations in GeoCas9(R1) had little effect on catalysis. In contrast, iGeoCas9 exhibited >6 fold faster DNA cleavage kinetics (0.37 min^-1^) compared to the wildtype or GeoCas9(R1) enzymes, underscoring the importance of WED domain mutations in promoting enzyme activity (Fig. 4C). Next, to test if the WED mutations affect DNA target sequence recognition, we re-designed the dsDNA substrate to include mismatches in the first two base pairs of the target region adjacent to a suboptimal 5’-N_4_GAAA-3’ PAM. Under single-turnover conditions, all three enzymes had similar initial rates for the cleavage of this substrate (0.21-0.26 min^-1^), indicating a five-fold increase in rate for both the wildtype GeoCas9 and GeoCas9(R1) enzymes, leading to activity levels similar to iGeoCas9 (Fig. 4C). This observation suggested that destabilization of the DNA double strands where target DNA and guide RNA base pairing begins compensates for the absence of iGeoCas9’s WED domain mutations. When tested with a linear substrate containing the native 5’-N_4_CAAA-3’ PAM, all three enzymes had similar cleavage kinetics (wildtype GeoCas9, 0.25 min^-1^; iGeoCas9, 0.30 min^-1^; GeoCas9(R1), 0.37 min^-1^) (Fig. 4C). Together, these data show that enzymes lacking the WED domain mutations, including wildtype GeoCas9 and GeoCas9(R1), require substrate destabilization by base pair mismatches to overcome the presence of a suboptimal PAM. These findings imply that the WED domain mutations supersede the role of the PAM in DNA target recognition by perhaps enhancing GeoCas9’s DNA untwisting capability.

**Figure 4.**
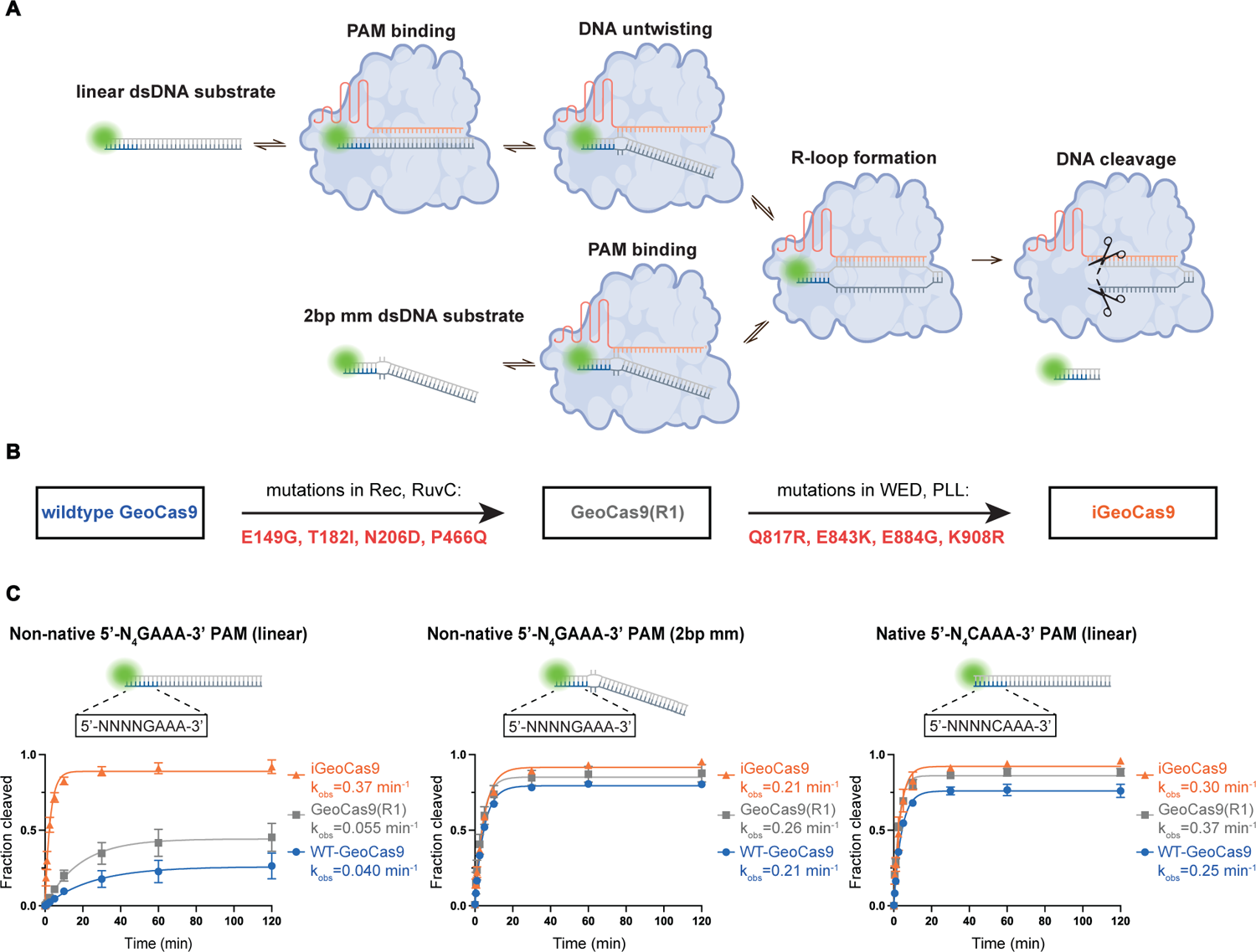
Thermodynamically unstable substrate mimics WED domain mutation effects on DNA melting. **(A)** Cleavage pathway for linear dsDNA substrates vs. two base pair mismatch (2 bp mm) dsDNA substrate. 60 nt DNA substrate is 5’ 6-FAM labeled (green). **(B)** The order in which mutations were introduced to create GeoCas9(R1) and iGeoCas9. Mutations are listed below the arrow, domains in which they are located are above the arrow. WED, wedge. PPL, phosphate lock loop. **(C)** In vitro dsDNA cleavage activity of wildtype (WT-)GeoCas9, iGeoCas9, and GeoCas9(R1) determined by denaturing PAGE (n = 3, mean ± SD). Substrate and PAM indicated above the graph. Fractions were collected at 0 sec, 30 sec, 1 min, 2.5 min, 5 min, 10 min, 30 min, 1 h, and 2 h. The k_obs_ for each Cas9 is listed in the sample legend.

### Faster R-loop formation by iGeoCas9 compensates for magnesium-restricted conditions

Wildtype GeoCas9 is ineffective for genome editing in mammalian cells, even when targeting genomic loci with optimal PAM sequences. In contrast, iGeoCas9 induces genome edits with >100-fold higher efficiency despite similar DNA cleavage kinetics compared to wildtype GeoCas9 using optimal substrates *in vitro*. One important difference between bacterial and mammalian cells is the availability of free magnesium ions, which can affect the formation of protein-nucleic acid complexes and Cas effector specificity^16, 17^. Bacterial cells and our biochemical assays have free magnesium concentrations of >1 mM^18^, whereas mammalian cells contain much lower levels of free magnesium ion of 0.1-1 mM^19,20^. To test the effect of magnesium ion concentration on enzyme activity, we determined DNA cleavage rates under single-turnover conditions for wildtype GeoCas9 and iGeoCas9 at 5-0.01 mM magnesium chloride concentrations. Using a 6-FAM-labeled substrate with a native 5’-N_4_CAAA-3’ PAM, we found that wildtype GeoCas9 and iGeoCas9 had similar single-turnover cleavage kinetics at 5 mM MgCl_2_ concentration.

However, a titration of magnesium concentrations from 1 mM to 0.1 mM resulted in a ∼17-fold decrease in wildtype GeoCas9-catalyzed DNA cleavage. Surprisingly, with a MgCl_2_ concentration as low as 0.01 mM, iGeoCas9 remains active with a k_obs_ of 0.043 min^-1^. Under the same conditions, wildtype GeoCas9 activity dropped to a k_obs_ below the detection limit (Fig. 5A and S5). These data suggest that iGeoCas9 is less dependent on magnesium, enabling it to maintain high activity in environments where free magnesium is less available such as mammalian cells.

**Figure 5.**
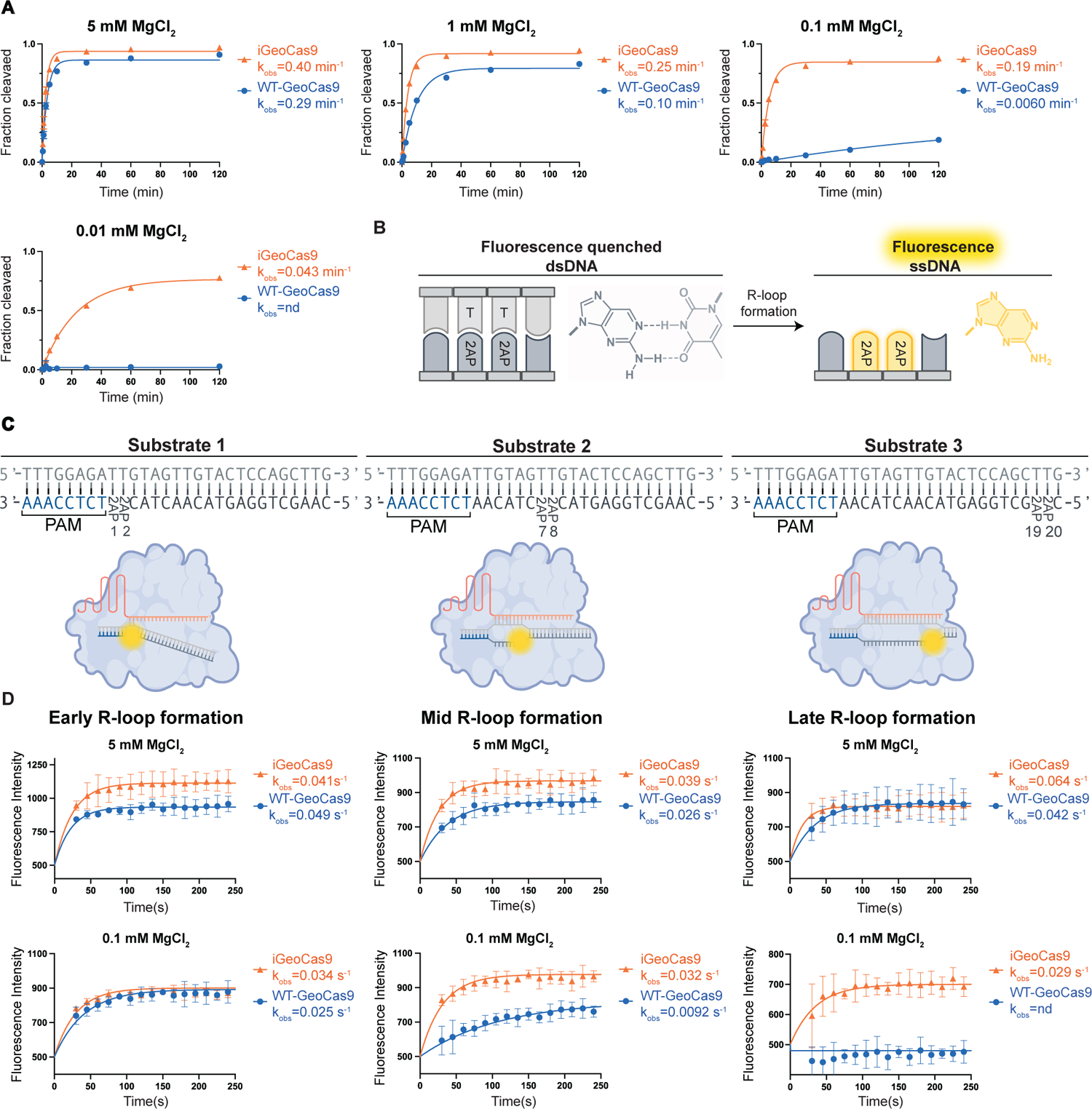
Reduced concentration of MgCl_2_ impacts wildtype GeoCas9 R-loop formation. **(A)** *In vitro* dsDNA cleavage activity of wildtype (WT-)GeoCas9 and iGeoCas9, determined by denaturing PAGE (n = 3, mean ± SD). MgCl**_2_** concentration indicated above the graph. Fractions were collected at 0 sec, 30 sec, 1 min, 2.5 min, 5 min, 10 min, 30 min, 1 h, and 2 h. The rate constants k_obs_ are listed in the sample legend. See also Figure S5. **(B)** Cartoon and chemical structure of 2-aminopurine (2AP) in the quenched and fluorescent states. Fluorescence indicated with yellow. T, Thymine. dsDNA, double stranded DNA. ssDNA, single stranded DNA. **(C)** Diagram of substrates (partial sequence) indicating positions of 2AP nucleotides measuring 5’ from the PAM (top). Drawing of the different stages of GeoCas9 R-loop formation with the three different substrates. Fluorescent 2APs are indicated in yellow (bottom). **(D)** 2AP fluorescence assays comparing catalytically inactivated wildtype GeoCas9 and iGeoCas9 R-loop kinetics (n =3, mean ± SD). Substrate 1, early R-loop formation. Substrate 2, mid R-loop formation. Substrate 3, late R-loop formation. MgCl_2_ concentrations indicated above the graph. The rate constant k_obs_ are listed in the sample legend.

The significant reduction of wildtype GeoCas9 activity at the low magnesium concentration found in human cells suggested a potential explanation for its poor genome editing activity in this cell type.

Furthermore, we wondered whether the mutations in iGeoCas9, which enable activity at very low magnesium ion concentrations *in vitro*, might have altered the rate-determining step of DNA cleavage. Since guide RNA strand invasion to form an R-loop with the DNA target sequence was previously determined to be the rate-limiting step for SpyCas9^17,21^, we tested whether there is a difference in the ability of wildtype GeoCas9 versus iGeoCas9 to form an R-loop under different magnesium ion conditions.

To investigate R-loop formation kinetics, we introduced 2-aminopurine (2AP) fluorescent nucleotides^21,22^ into the non-target strand of the dsDNA substrate containing an optimal PAM 5’-N_4_CAAA-3’ sequence. In the dsDNA helix, 2AP fluorescence is quenched through base stacking interactions with neighboring nucleotides. The unwinding and destacking of dsDNA by Cas9 results in 2AP fluorescence (Fig. 5B)^23^. To determine R-loop formation kinetics at different stages, dsDNA substrates were designed to contain tandem 2AP markers located at varying distances from the PAM. Three substrates with 2APs at positions 1&2, 7&8 or 19&20 of the target region were used to determine the early, mid, and late stages of R-loop formation kinetics, respectively (Fig. 5C). Consistent with prior measurements, no major differences in kinetics were observed between wildtype GeoCas9 and iGeoCas9 when testing the three substrates in the presence of 5 mM MgCl_2_ (Fig. 5D). Interestingly, when the MgCl_2_ concentration was reduced to 0.1 mM, reactions using substrate 1 with 2APs at positions 1&2 showed similar early R-loop formation kinetics for both wildtype GeoCas9 and iGeoCas9, suggesting there is no major difference in the initial DNA interrogation step (dsDNA untwisting) when using an optimal PAM 5’-N_4_CAAA-3’ (Fig. 5D). However, a 3.5-fold difference was observed for the kinetics of mid R-loop formation (2AP at positions 7&8) when comparing wildtype GeoCas9 and iGeoCas9. More surprisingly, the kinetic analyses of late R-loop formation (2AP at positions 19&20) established that wildtype GeoCas9 is exceptionally slow at dsDNA unwinding under MgCl_2_-restricted conditions, while iGeoCas9 maintained fast kinetics for complete R-loop formation with a kinetic constant (0.029 s^-1^) comparable to that of early R-loop (0.034 s^-1^) (Fig. 5D).

R-loop kinetic measurements show that in the presence of high MgCl_2_ concentrations (e.g., 5 mM) early, mid and late R-loop formation were similar for wildtype GeoCas9 and iGeoCas9. This suggests that under these conditions, R-loop formation is not the rate-determining step. Under conditions of low MgCl_2_ concentration (e.g., 0.1 mM), mid and late R-loop formation by wildtype GeoCas9 were slow, suggesting that R-loop formation has become rate-determining. In contrast, iGeoCas9 quickly progressed through all stages of R-loop formation at low MgCl_2_. Together, these results establish that the mutations in iGeoCas9 substantially improve DNA unwinding capability in the mid and late stages of R-loop formation, enabling sustained enzyme activity under low magnesium ion conditions.

### Mutations accelerating R-loop formation are transferable to another genome editor

Having established that iGeoCas9 WED domain mutations are likely to promote genome editing efficiency through expedited R-loop formation and reduced PAM specificity, we wondered whether these biochemical insights might enable rational engineering of other Cas9 proteins. Nme2Cas9^24,25^, with ∼38% sequence identity to GeoCas9, was selected as a candidate protein for engineering. Similar to iGeoCas9, WED domain mutations were introduced into Nme2Cas9 to generate two new improved versions of Nme2Cas9. Nme2Cas9(v1) contains E868K, K870R and K929R mutations to allow for potential new DNA target strand interactions, and D873A and D911G mutations that could mimic the E884G mutation in iGeoCas9 (Fig. 6A and S6A). Two additional mutations, E932K and D844G, expected to interact with the non-target DNA strand and alter protein charge respectively, were further introduced to generate iNme2Cas9 (Figs. 6A, S6B). The genome editing activities of wildtype Nme2Cas9, Nme2Cas9(v1) and iNme2Cas9 were evaluated using an enhanced green fluorescent protein (EGFP) knock-down assay in HEK293T cells. Plasmids encoding Nme2Cas9 proteins together with sgRNAs were transfected to HEK293T cells, which were analyzed by flow cytometry to give editing efficiencies (Fig. 6B). Six guide RNAs, guide 1 to 6, were designed to target the EGFP transgene using various PAM sequences. 5’-N_4_CC-3’ was previously identified as the optimal PAM sequence for wildtype Nme2Cas9^23^. Guides 1 and 2 were designed to use the native 5’-N_4_CC-3’ PAM, and the corresponding editing tests to disrupt EGFP revealed substantial improvement in the editing efficiency from wildtype to Nme2Cas9(v1) and iNme2Cas9 (Figs. 6C, S6C). Based on our demonstration that the WED domain mutations in iGeoCas9 led to relaxed PAM recognition, we expected to see an expansion in PAM sequences with our engineered Nme2Cas9 variants. Therefore, guides 3-6, targeting the EGFP gene with non-native PAMs, 5’-N_4_CT-3’, 5’-N_4_TC-3’, 5’-N_4_CA-3’, and 5’-N_4_TT-3’, respectively, were also included in our editing test. As expected, our engineered iNme2Cas9 was able to give up to 50% EGFP knock-down efficiency with guides 3-6, while the wildtype Nme2Cas9 barely induced any editing activity (Fig. 6C). Overall, our iNme2Cas9 showed up to >100-fold improvement in genome editing activity compared to the wildtype enzyme.

**Figure 6.**
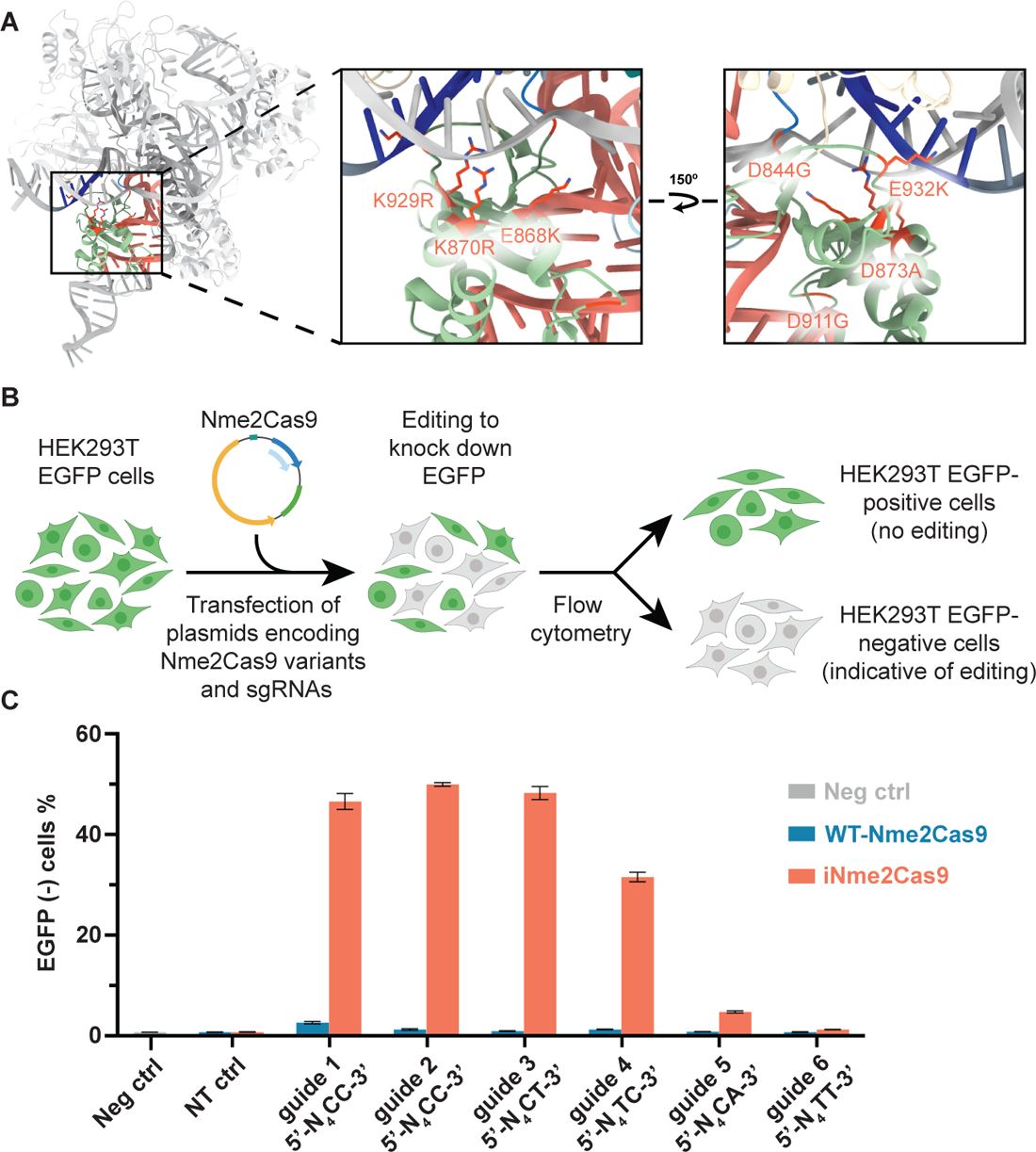
WED domain mutations greatly enhance genome editing activities of Nme2Cas9. **(A)** Model of Nme2Cas9 (PDB:6JE3)^25^ in which all seven rationally engineered mutations are represented in red. Rotamers were chosen to demonstrate potential DNA interactions. **(B)** Workflow for EGFP-knock-down assay in HEK293T cells. Successful editing indicated by a loss of EGFP signal. **(C)** HEK293T cell editing by wildtype (WT-) Nme2Cas9 and iNme2Cas9, and 6 different guides. Neg, no treatment control; NT, non-targeting guide control; See also Figure S6.

Nme2Cas9 has been previously engineered using a phage-assisted continuous evolution (PACE) system by the Liu lab^26^. We were thus interested in comparing our engineered iNme2Cas9 to the reported variants. We selected two nuclease-reactivated mutants from the study, Nme2Cas9(C-NR) and Nme2Cas9(T-NR), which recognize C- and T-based PAM sequences, respectively. Using the same EGFP knock-down assay under dose-limiting conditions, we were delighted to observe that our rationally designed iNme2Cas9 outperformed the previously engineered mutants using five guide RNAs and single-pyrimidine PAMs (Fig. S6D). These results have further reinforced that the WED domain mutations are responsible for accelerating R-loop formation under magnesium-restricted conditions, typical of mammalian cell environments, which, however, may not be preferentially selected out using bacterial-based evolution systems like PACE. Taking all these together, we believe our rational engineering of Nme2Cas9 supports that optimizing Cas9 WED domain can lead to robust genome editors with expanded PAM compatibility and improved editing efficiency.

## DISCUSSION

CRISPR-Cas9-based technology has diverse applications across the life sciences^5,27^. In particular, Cas9 has the therapeutic potential to treat a wide range of genetic diseases^28,29^. To expand its utility, the basic functions of CRISPR-Cas9 have been extended using protein engineering^30^. For example, engineered SpyCas9 enzymes can accommodate a range of non-NGG PAM sequences, enabling them to target a wider range of genomic sites^12,13,31^. Cas9 proteins have also been rationally optimized to have higher fidelity of DNA cleavage, leading to reduced off-target editing^32–34^. However, protein engineering has demonstrated little success in improving Cas9’s basic genome editing activities^34^. This may be because we lack a proper understanding of the essential relationship between Cas9 structure and its genome editing activities. A separate study demonstrated that engineering a thermostable type II-C Cas9 from *Geobacillus stearothermophilus* (GeoCas9) can produce a robust genome editor, iGeoCas9, with substantially improved editing efficiency in cultured mammalian cells and animal tissues^9^. Biochemical and structural comparison of wildtype GeoCas9 versus iGeoCas9 as reported here provides insights into the structure-function relationship of Cas9 and establishes general principles for engineering Cas9’s genome editing activities.

Our work reveals the pivotal and unanticipated role that the WED domain plays in regulating type II-C Cas9’s genome editing activities. We found that WED domain mutations in iGeoCas9 establish new interactions with the DNA target strand and non-target strand backbones in the PAM region of the dsDNA target. These interactions enhance R-loop formation, the DNA interrogation step that involves unwinding the target dsDNA sequence to enable RNA-DNA duplex formation. R-loop formation is the rate-limiting step in Cas9-catalyzed DNA cleavage^16,21^, and slow R-loop formation kinetics likely limits GeoCas9’s editing activities in mammalian cells. We found that the evolved editor, iGeoCas9, was able to overcome the slow R-loop kinetics observed with wildtype GeoCas9 in low magnesium environments. This effect could be due to the observed increased interactions between the WED domain and target dsDNA, leading to a dramatically accelerated R-loop formation process. This makes R-loop formation no longer the rate-limiting step in catalysis and also imposes a much less stringent requirement on PAM recognition for target binding.

Having evolved as powerful immune defense machinery in prokaryotic organisms, heterologous biological environments may alter or restrict the targeted endonucleolytic function of CRISPR-Cas enzymes. A better understanding of the mechanisms and constraints of CRISPR enzymes, followed by the development of general engineering strategies to attenuate these constraints, can help establish robust and effective genome editing tools. Our study illustrates that enhanced binding between the Cas9-guide RNA enzymatic complex and the target dsDNA corresponds to increased genome editing efficiency. In particular, the three WED domain mutations in iGeoCas9 create new DNA TS and NTS backbone interactions and alter the electrostatic environment of the protein-DNA interface, accelerating R-loop formation under magnesium-restricted conditions. Similar engineering of the related Nme2Cas9 protein generated a mutant editor showing remarkable improvement for mammalian cell genome editing. Several groups have reported an engineering strategy that involves introducing cationic residues to enhance target DNA binding, thereby improving the function of CRISPR-Cas or relevant proteins^36–38^.

However, this strategy requires intensive mutational screening at residues surrounding target DNA or even arginine scanning throughout the entire protein. Our findings highlight the WED domain as a previously unappreciated regulator of the mechanisms that drive Cas9’s genome editing behaviors, particularly in mammalian cells with low free magnesium content. The WED domain, functioning as a bridge between the sgRNA (or crRNA) scaffold and the upstream dsDNA region of the target DNA, can be identified in CRISPR-Cas9, Cas12, and their relevant proteins, including IscB^39^, TnpB^40^, and Fanzor^41^ proteins. Despite different structural organizations of the WED domain among the diverse systems, we assume that engineering of the WED domain (or other related regions with similar structural functions, e.g., PI domain) can possibly be applied to these systems based on the simple principle to enhance the binding to target DNA. Preliminary findings with the state-of-the-art genome editor, type II-A SpyCas9, albeit featuring different C-terminal domain organization (WED + PI) compared to type II-C Cas9s, show that it can also be potentially engineered with improved editing efficiency (Fig. S6). Therefore, we believe focusing engineering efforts on the dsDNA-binding regions of CRISPR-Cas proteins can be a streamlined approach that holds promise for the development of more effective genome editing systems and would lead to more robust CRISPR-based therapeutics.

## ACKNOWLEDGEMENTS

We thank members of the Doudna lab and the Innovative Genomics Institute for helpful discussions. We also thank Daniel B. Toso and Ravindra Thakkar at the University of California Berkeley QB3 Cal-Cryo Facility for their support with operating the Talos Arctica and Titan Krios G2. We would also like to acknowledge Ms. Netravathi Krishnappa (NGS Core Operations Manager and Sequencing Specialist, Center for Translational Genomics, Innovative Genomics Institute, UC Berkeley) for NGS. This work was supported by the Life Sciences Research Foundation (K.C.); the National Institutes of Health (AI17111, A.R.E); the CIRM Training Program (EDUC4-12790). Figure cartoons generated with biorender.com. J.A.D. is an investigator of the Howard Hughes Medical Institute (HHMI) and is funded by HHMI.

## AUTHOR CONTRIBUTIONS

Conceptualization: A.R.E, K.C., and J.A.D.; experimental studies: A.R.E, K.C., K.M.S, O.T.T., E.E.D, B.W.T., and B.X.; data analysis: A.R.E, K.C., K.M.S, O.T.T., E.E.D, B.W.T., and M.I.T.; supervision: J.A.D.; manuscript writing: A.R.E., K.C. and J.A.D. with input from all authors.

## DECLARATION OF INTERESTS

The Regents of the University of California have patents issued and pending for CRISPR technologies on which the authors are inventors. J.A.D. is a cofounder of Caribou Biosciences, Editas Medicine, Scribe Therapeutics, Intellia Therapeutics, and Mammoth Biosciences. J.A.D. is a scientific advisory board member or consultant for Vertex, Caribou Biosciences, Intellia Therapeutics, Scribe Therapeutics, Mammoth Biosciences, Algen Biotechnologies, Felix Biosciences, The Column Group, and Inari. J.A.D. is Chief Science Advisor to Sixth Street, a Director at Altos, Johnson & Johnson and Tempus, and she has research projects sponsored by Apple Tree Partners and Roche.

## METHOD DETAILS

### GeoCas9 protein expression and purification

Protein expression and purification were performed previously as described with some modifications^7^. Briefly, *E. coli* BL21 DE3 cells were transformed with Cas9 expression plasmids in 2xYT medium supplemented with 100 ug/ml ampicillin and grown to an OD between 0.6 to 0.8. Cultures were cooled on ice for 30 min then supplemented with 0.5 mM isopropyl b-D-thiogalactoside (IPTG) and grown overnight at 16℃. Cells were harvested by centrifugation, gently resuspended in lysis buffer (50 mM Tris-HCl, pH 7.5, 20 mM imidazole, 0.5 mM TCEP, 500 mM NaCl, 1 mM PMSF), and lysed by sonication. Clarified lysate was incubated with Ni-NTA resin pre-equilibrated with wash buffer (50 mM Tris-HCl, pH 7.5, 20 mM imidazole, 0.5 mM TCEP, 500 mM NaCl) for 1 hour. Protein bound resin was washed 3 times and eluted with Ni-NTA elution buffer (50 mM Tris-HCl, pH 7.5, 500 mM imidazole, 0.5 mM TCEP, 500 mM NaCl).

Pierce Human Rhinovirus 3C Protease was added to the elution for cleavage and dialyzed overnight in a 10,000 MWCO dialysis cassette with dialysis buffer (20 mM HEPES pH 7.5, 150 mM KCl, 10% glycerol, and 1 mM TCEP) overnight at 4℃. Proteins were filtered with a 0.22 uM filter unit (Millex GP), injected into a pre-equilibrated MBPTrap HP column (Cytiva) and washed buffer A (20 mM HEPES pH 7.5, 150 mM KCl, 10% glycerol, and 1 mM TCEP) until 260 and 280 absorbance readings reached UV baseline. Proteins were eluted with a linear gradient of KCl concentrated with a 30,000 MWCO concentrator (Millipore Sigma), and loaded onto a Superdex 200 Increase 10/300 GL and purified using gel filtration buffer (20 mM HEPES, pH 7.5, 150 mM KCl, 10% glycerol, 1 mM TCEP). Peak fractions containing Cas9 were quantified using a NanoDrop 8000 Spectrophotometer (Thermo Scientific) and flash frozen and stored at −80°C.

### Nucleic acid preparation

DNA substrates and sgRNA was ordered from Integrated DNA Technologies (IDT). Substrates were purified in house using a 12% urea-PAGE gel. The band containing the substrate was excised, gentle ground, and incubated in 1:10 volume of 3M sodium acetate overnight at 4°C. Samples were 0.22 um vacuum filtered and concentrated with a 3 kDa MWCO centrifugal filter (MilliporeSigma). Samples were ethanol precipitated with greater than 2.5 times the volume of 100% ethanol at −80°C for 2 hours. Samples were pelleted using centrifugation, washed twice with 80% ethanol, and resuspended in molecular grade water. Samples were stored at −80°C for later use.

### Cryo-EM grid preparation and data collection

Wildtype GeoCas9 complexes were frozen on UltrAuFoil R 1.2/1.3 (Electron Microscopy Sciences product num. Q350AR13A) grids using FEI Vitrobot Mark IV set to 8°C with 100% humidity. Grids were glow discharged at 15 mA for 25 s using Pelco easyGLOW. 4 µl of sample was applied to the grids and blotted for 4 s with blot force 8. Micrographs for the wildtype complex were collected on Talos Arctica operated at 200 kV and 36,000x nominal magnification (1.14 Å pixel size) in super resolution mode (0.57 Å pixel size) on K3 Direct Electron Detector in CDS mode. Movies were collected using SerialEM version 4.0.10 with 50 e/Å^2^ final dose. Micrographs for the mutant complex were collected on Titan Krios G2 operated at 300 kV and 81,000x nominal magnification (0.93 Å pixel size) in super resolution mode (0.465 Å pixel size) on K3 Direct Electron Detector in CDS mode. Movies were collected using SerialEM version 4.0.19 with 50 e/Å^2^ final dose.

### Cryo-EM data processing

2,767 movies with the wildtype GeoCas9 complex were collected with the defocus range −0.8 to −2. Movies were processed in CryoSPARC software (Structura Biotechnology) version 4.2.1. Movies were corrected for beam-induced motion with patch motion and CTF parameters were calculated with patch CTF. After manual curation 2,353 micrographs remained for further analysis. Particle picking was optimized using blob, template and Topaz picking resulting in the extraction of 1,026,723 particles^42^. They were subjected to 2D classification, and the best classes were selected leaving 949,346 particles.

Particles were re-extracted with recentering and subjected to *ab initio* reconstruction into two classes. Only one class resembled Cas9 complex and contained 645,337 particles. Particles from this class were re-extracted with recentering again. This *ab initio* class was then refined with re-extracted particles using non-uniform refinement resulting in an anisotropic map. Particles were again re-extracted with recentering and classified using 3D classification into 6 classes in the simple mode. Each class was refined using non-uniform refinement^43^. After refinement only one class was isotropic, had the best completeness and reached the highest resolution of 3.17 Å. The best class contained 117,726 particles and served as a basis for model building.

For the iGeoCas9 complex 7,849 movies were collected with defocus range −0.8 to −2. Data was processed in an analogical way as the wildtype complex. After beam-induced motion correction, ctf estimation and manual curation 5,731 micrographs remained. Particles were picked with the blob picker. 6,498,580 extracted particles were subjected to two rounds of 2D classification and class selection. 1,840,865 selected particles were re-extracted with recentering and used for *ab initio* map reconstruction into two classes. 1,368,998 particles from the good class were re-extracted with recentering again and used for refinement of the good class with non-uniform refinement, which resulted in an anisotropic map. Particles from the refinement job were used for the further classification using 3d classification into 6 classes in the simple mode. Each class was refined with non-uniform refinement and the class which reached the highest resolution of 2.63 Å was also isotropic and served as a final map for model building^43^. The final map was obtained from 228,251 particles.

### Model building

The initial model of wildtype GeoCas9 was generated by ModelAngelo v1.0.1 and was built to wildtype GeoCas9 map that was resampled to pixel size 0.93, and box size 256. Missing residues in the areas of reduced quality density were filled manually with corresponding regions from the Colabfold v1.4.0 model. The density for the following domains was missing in the sharp map and was omitted in the final wildtype GeoCas9 model: Rec-I, residues 134 to 140; HNH, residues 524-665; RuvC-III, residues 749-755; PI residues, 1067-1087. Nucleic acids were built based on Nme2Cas9-sgRNA-dsDNA (PDB:6JE3)^25^ as an initial model. The density for the following nucleic acids was missing in the sharp map and was omitted in the final wildtype model: sgRNA (5’-3’) nucleotides 71-75, 106-128, 136-139; TS (5’-3’) 1-4, 49-51; and NTS (5’-3’) nucleotides 1-30, and 51. Wildtype GeoCas9 model went through several rounds of real space refinement in Phenix version 1.19.2-4158 and manual geometry improvement in Coot version 0.9.8.7 resulting in a final model^44,45,46^.

The final wildtype model served as the initial model for iGeoCas9, with the introduction of the iGeoCas9 mutations. The iGeoCas9 map was resampled to pixel size 0.93 A, and box size 256. The density for the additional nucleic acids was missing in the sharp map and was omitted in the final iGeoCas9 model:TS (5’-3’) 5 and 45-48; and NTS (5’-3’) 47-50. Model was refined in Phenix with real space refinement. Geometry was improved manually in Coot and Phenix refinement was repeated to obtain the final iGeoCas9 model^44,45,46^.

### In vitro cleavage assays

For linear dsDNA cleavage assays target strands were labeled on the 5’ end with 6-FAM for visualization. sgRNA were annealed in 1 x annealing buffer (10 mM Tris, pH 7.5, 20 mM KCl, 1.5 mM MgCl_2_) by heating at 80°C for 2 minutes then placing it on ice. GeoCas9 was incubated with 1.3 x sgRNA in 1 x cleavage buffer (20 mM Tris-HCl, pH7.5, 100 mM KCl, 5 mM MgCl_2,_ 1 mM TCEP, 5% (w/v) glycerol) at room temperature for 30 min followed by 5 min at 37°C for RNP formation. For reduced magnesium cleavage assays, final magnesium concentration for the 1 x cleavage buffer are as indicated in the results. The final concentration of GeoCas9 RNP was 100 nM and FAM-labeled substrates was 20 nM. Cleavage reactions were initiated by mixing GeoCas9 RNP and FAM-labeled substrate on a 37°C thermoblock. Sample fractions were collected at 0, 1, 2.5, 5, 10, 30, 60, 120 minutes and mixed with 2 x quench buffer (94% (v/v) formamide, 30 mM EDTA, 400 mg/mL heparin, 0.2% SDS, and 0.025% (w/v) bromophenol blue) to stop the reaction. Quenched samples were heated at 95°C for 2 minutes and resolved on a denaturing PAGE gel (12% acrylamide:bis-acrylamide 19:1, 7 M urea, 1X TBE). Gels were visualized using a phosphorimager (Amersham Typhoon, GE Healthcare) and quantified using Cytiva ImageQuantTL 10.1.

For time course experiments, kinetic analyses of substrates were performed in triplicate. Cleaved fractions were determined by dividing the cleaved target strand density by the total density (cleaved + uncleaved) for each sample. Quantified data was fit to a curve generated by the formula Y=Y_max_*(1-exp(-k*t)) where Y_max_ is the pre-exponential factor, k is the rate constant kobs, and t is the reaction time in minutes. If Y_max_ ≤ 0.05, k_obs_ is not determined “nd”.

### PAM depletion assays

PAM depletion assays were performed with purified iGeoCas9 and WT GeoCas9. RNPs were formed as instructed above (see methods *in vitro* cleavage assays) using 1.25 x the amount of sgRNA to Cas9. Samples included both targeting and non-targeting guides for iGeoCas9 and WT Cas9. An “untreated” control was processed alongside in which no RNP was added to the reaction. Libraries were constructed with PAM 5’-“TTTTNNNN”-3 downstream of the spacer on the non-target strand, where (N) represents randomized nucleotides. A 206 bp library fragment was generated using overlap PCR and included NexteraXT adapter overhang sequences (*Illumina 16S Metagenomic Sequencing Library Preparation* protocol, Part # 15044223 Rev. B) flanking the target region 20 bp (3’) downstream and 98 bp (5’) upstream of the target sequence. 500 nM of Cas9 RNP was incubated with a 100 nM library for 2 hours at 37℃. Library DNA was purified using 0.8 x Ampure beads (Beckman Coulter) and washed three times with 80% ethanol. DNA bound beads were air dried and resuspended in 10 mM Tris-Cl, pH 8.5. Non cleaved library fragments containing both adapter sites were preferentially amplified using Q5 High-Fidelity DNA Polymerase and indexed with NextraXT adapters using 12 cycles of PCR, producing a final library size of 279 bp. PCR products were cleaned with Ampure beads as instructed previously. Libraries were quantified using qPCR (Kapa Biosystems) and normalized, pooled to 10 nM, and submitted to the Innovative Genomics Institute (IGI) Next Generation Sequencing Core for 100 x sequencing coverage on the NextSeq.

### NGS analysis for PAM depletion assays

PAM specificity was characterized from FASTQs using a custom Python script. Briefly, regular expressions were used to extract four-nucleotide PAMs. PAM frequencies were calculated and normalized to the sequencing depth of each sample, then the log2-fold-change relative to frequencies in the untreated library was determined. Significantly depleted PAMs were defined as those exceeding the 99.9999% confidence interval for maximum log2-fold-change depletion in the non-targeting samples. Sequence logos were generated from significant PAMs with Logomaker version 0.8.

### Monitoring of R-loop formation via 2-aminopurine

DNA duplexes were designed so that 2-aminopurine (2AP) is on the NTS^23^. There were 3 substrates total with 2 sequential 2AP molecules in positions 1&2, 7&8, 19&20 from the PAM sequence. DNA was annealed at a 1:1.2 molar ratio in (50 mM Tris, pH 7.5, 100 mM NaCl) by heating to 95°C for 3 minutes and then cooling to room temperature over 45 minutes. To prepare the ternary complex, sgRNA was heated to 95°C for 1 min in ME buffer (10 mM MOPs, pH 6.5 and 1 mM EDTA). To the sgRNA was added 1X reaction buffer (20 mM Tris, pH 7.5; 100 mM KCl, 5% glycerol, with 0.1 mM MgCl_2_ or 5 mM MgCl_2_) before adding Cas9 to assemble sgRNA/GeoCas9 at a 1:1.25 molar ratio in 1X reaction buffer. The GeoCas9/sgRNA mixture was incubated at room temperature for 30 minutes. The reaction was initiated by the addition of the GeoCas9/sgRNA mixture to duplex DNA containing 2-aminopurine at a final molar ratio of 5:1 in 1X reaction buffer.

### HEK293T EGFP editing assay

HEK293T EGFP reporter cells were seeded in 96-well plates (20k cell/well) and transfected 24 h later at ∼70% confluency according to the manufacturer’s protocol with lipofectamine 3000 (Life Technologies) and 100 ng (50 ng used for dose-limiting conditions) of plasmid DNA encoding the wildtype or engineered Nme2Cas9 and sgRNA. 48 h post-transfection, HEK293T EGFP reporter cells were subjected to selection with 2 μg/mL puromycin in cell culture media for 48 h. Cell culture media was refreshed to exclude puromycin. Cells were collected in 96-well round bottom plates after trypsinization for flow analysis on an Attune NxT Flow Cytometer with an autosampler.

## REFERENCES

1. Sorek, R., Lawrence, C.M., and Wiedenheft, B. (2013). CRISPR-mediated adaptive immune systems in bacteria and archaea. Annu. Rev. Biochem. 82, 237–266. 10.1146/annurev-biochem-072911-172315

2. Jiang, F., and Doudna, J.A. (2017). CRISPR–Cas9 structures and mechanisms. Annu. Rev. Biophys. 46, 505–529. 10.1146/annurev-biophys-062215-010822

3. Jinek, M., Chylinski, K., Fonfara, I., Hauer, M., Doudna, J.A., and Charpentier, E. (2012). A programmable dual-RNA–guided DNA endonuclease in adaptive bacterial immunity. Science 337, 816– 821. 10.1126/science.1225829

4. Cong, L., Ran, F.A., Cox, D., Lin, S., Barretto, R., Habib, N., Hsu, P.D., Wu, X., Jiang, W., Marraffini, L.A., and Zhang, F. (2013). Multiplex genome engineering using CRISPR/Cas systems. Science 339, 819–823. 10.1126/science.1231143

5. Wang, J. Y., and Doudna, J. A. (2023). CRISPR technology: A decade of genome editing is only the beginning. Science 379, eadd8643. https://doi/10.1126/science.add8643

6. Camperi, J., Moshref, M., Dai, L., and Lee, H. Y. (2022). Physicochemical and functional characterization of differential CRISPR-Cas9 ribonucleoprotein complexes. Anal. Chem. 94, 1432–1440. https://pubs.acs.org/doi/10.1021/acs.analchem.1c04795

7. Harrington, L. B., Staahl, B. T., Chen, J. S., Ma, E., Kyrpides, N. C., and Doudna, J. A. (2017). A thermostable Cas9 with increased lifetime in human plasma. Nat. Comm. 8, 1424. 10.1038/s41467-017-01408-4

8. Mougiakos, I., Mohanraju, P., Bosma, E. F., Vrouwe, V., Finger Bou, M., Naduthodi, M. I., Gussak, A., Brinkman, R. B., and Van Kranenburg, R. (2017). Characterizing a thermostable Cas9 for bacterial genome editing and silencing. Nat. Comm. 8, 1647. 10.1038/s41467-017-01591-4

9. Chen, K., Han, H., Zhao, S., Xu, B., Yin, B., Trinidad, M., Burgstone, B. W., Murthy, N., and Doudna, J. A. (2023). Lung and liver editing using lipid nanoparticle delivery of a stable CRISPR-Cas9. BioRxiv. 10.1101/2023.11.15.566339

10. Anders, C., Niewoehner, O., Duerst, A., and Jinek, M. (2014). Structural basis of PAM-dependent target DNA recognition by the Cas9 endonuclease. Nature 513, 569–573. 10.1038/nature13579

11. Karvelis, T., Gasiunas, G., Young, J., Bigelyte, G., Silanskas, A., Cigan, M., and Siksnys, V. (2015). Rapid characterization of CRISPR-Cas9 protospacer adjacent motif sequence elements. Genome Biol. 16, 253. 10.1186/s13059-015-0818-7

12. Hu, J. H., Miller, S. M., Geurts, M. H., Tang, W., Chen, L., Sun, N., Zeina, C. M., Gao, X., Rees, H. A., Lin, Z., and Liu, D. R. (2018). Evolved Cas9 variants with broad PAM compatibility and high DNA specificity. Nature 556, 57–63. 10.1038/nature26155

13. Walton, R. T., Christie, K. A., Whittaker, M. N., and Kleinstiver, B. P. (2020). Unconstrained genome targeting with near-PAMless engineered CRISPR-Cas9 variants. Science 368, 290–296. 10.1126/science.aba8853

14. Zhang, Y., Zhang, H., Xu, X., Wang, Yujue, Chen, W., Wang, Yannan, Wu, Z., Tang, N., Wang, Yu, Zhao, S., Gan, J., Ji, Q. (2020). Catalytic-state structure and engineering of Streptococcus thermophilus Cas9. Nat. Catal. 3, 813–823. 10.1038/s41929-020-00506-9

15. Ma, E., Harrington, L. B., O’Connell, M. R., Zhou, K., and Doudna, J. A. (2015). Single-stranded DNA cleavage by divergent CRISPR-Cas9 enzymes. Mol. Cell 60, 398–407. 10.1016/j.molcel.2015.10.030

16. Raper, A. T., Stephenson, A. A., and Suo, Z. (2018). Functional insights revealed by the kinetic mechanism of CRISPR/Cas9. J. Am. Chem. Soc. 140, 2971–2984. https://pubs.acs.org/doi/10.1021/jacs.7b13047

17. Nguyen, G.T., Schelling, M.A., Buscher, K.A., Sritharan, A., Sashital, D.G. (2023). CRISPR-Cas12a exhibits metal-dependent specificity switching. bioRxiv 2023.11.29.569287. 10.1101/2023.11.29.569287

18. Groisman, E. A., Hollands, K., Kriner, M. A., Lee, J., Park, Y., and Pontes, M. H. (2013). Bacterial Mg^2+^ homeostasis, transport, and virulence. Annu. Rev. Genet. 47, 625–646. 10.1146/annurev-genet-051313-051025

19. Fatholahi, M., LaNoue, K., Romani, A., and Scarpa, A. (2000). Relationship between total and free cellular Mg^2+^ during metabolic stimulation of rat cardiac myocytes and perfused hearts. Arch. Biochem. Biophys. 374, 395–401. 10.1006/abbi.1999.1619

20. Raju, B., Murphy, E., Levy, L. A., Hall, R. D., and London, R. E. (1989). A fluorescent indicator for measuring cytosolic free magnesium. Am. J. Physiol. Cell Physiol. 256, 540–548. 10.1152/ajpcell.1989.256.3.C540

21. Gong, S., Yu, H. H., Johnson, K. A., and Taylor, D. W. (2018). DNA unwinding is the primary determinant of CRISPR-Cas9 activity. Cell Rep. 22, 359–371. 10.1016/j.celrep.2017.12.041

22. Jean, J. M., and Hall, K. B. (2001). 2-Aminopurine fluorescence quenching and lifetimes: Role of base stacking. Proc. Natl. Acad. Sci. U.S.A. 98, 37–41. 10.1073/pnas.98.1.37

23. Li, Y., Liu, Y., Singh, J., Tangprasertchai, N. S., Trivedi, R., Fang, Y., and Qin, P. Z. (2022). Site-specific labeling reveals Cas9 induces partial unwinding without RNA/DNA pairing in sequences distal to the PAM. CRISPR J. 5, 341–352. 10.1089/crispr.2021.0100

24. Edraki, A., Mir, A., Ibraheim, R., Gainetdinov, I., Yoon, Y., Song, C., Cao, Y., Gallant, J., Xue, W., Rivera-Pérez, J. A., and Sontheimer, E. J. (2019). A compact, high-accuracy Cas9 with a dinucleotide PAM for *in vivo* genome editing. Mol. Cell 73, 714–726. 10.1016/j.molcel.2018.12.003

25. Sun, W., Yang, J., Cheng, Z., Amrani, N., Liu, C., Wang, K., Ibraheim, R., Edraki, A., Huang, X., Wang, M., Wang, J., Liu, L., Sheng, G., Yang, Y., Lou, J., Sontheimer, E. J., and Wang, Y. (2019). Structures of *Neisseria meningitidis* Cas9 complexes in catalytically poised and anti-CRISPR-inhibited states. Mol. Cell 76, 938–952. 10.1016/j.molcel.2019.09.025

26. Huang, T. P., Heins, Z. J., Miller, S. M., Wong, B. G., Balivada, P. A., Wang, T., Khalil, A. S., and Liu, D. R. (2023). High-throughput continuous evolution of compact Cas9 variants targeting single-nucleotide-pyrimidine PAMs. Nat. Biotech. 41, 96–107. 10.1038/s41587-022-01410-2

27. Gersbach, C. A. (2019). The next generation of CRISPR–Cas technologies and applications. Nat. Rev. Mol. Cell Biol. 20, 490–507. 10.1038/s41580-019-0131-5

28. Li, T., Yang, Y., Qi, H., Cui, W., Zhang, L., Fu, X., He, X., Liu, M., Li, P., and Yu, T. (2023). CRISPR/Cas9 therapeutics: Progress and prospects. Signal Transduct. Target. Ther. 8, 36. 10.1038/s41392-023-01309-7

29. Sharma, G., Sharma, A. R., Bhattacharya, M., Lee, S., and Chakraborty, C. (2021). CRISPR-Cas9: A preclinical and clinical perspective for the treatment of human diseases. Mol. Ther. 29, 571–586. 10.1016/j.ymthe.2020.09.028

30. Liu, R., Liang, L., Freed, E. F., and Gill, R. T. (2021). Directed evolution of CRISPR/Cas systems for precise gene editing. Trends Biotechnol. 39, 262–273. https://www.cell.com/trends/biotechnology/fulltext/S0167-7799(20)30200-6.

31. Nishimasu, H., Shi, X., Ishiguro, S., Gao, L., Hirano, S., Okazaki, S., Noda, T., Abudayyeh, O. O., Gootenberg, J. S., Mori, H., Oura, S., Holmes, B., Tanaka, M., Seki, M., Hirano, H., Aburatani, H., Ishitani, R., Ikawa, M., Yachie, N., Zhang, F., Nureki, O. (2018). Engineered CRISPR-Cas9 nuclease with expanded targeting space. Science 361, 1259–1262 10.1126/science.aas9129

32. Kleinstiver, B. P., Pattanayak, V., Prew, M. S., Tsai, S. Q., Nguyen, N. T., Zheng, Z., and Joung, J. K. (2016). High-fidelity CRISPR–Cas9 nucleases with no detectable genome-wide off-target effects. Nature 529, 490–495. 10.1038/nature16526

33. Chen, J. S., Dagdas, Y. S., Kleinstiver, B. P., Welch, M. M., Sousa, A. A., Harrington, L. B., Sternberg, S. H., Joung, J. K., Yildiz, A., and Doudna, J. A. (2017). Enhanced proofreading governs CRISPR–Cas9 targeting accuracy. Nature 550, 407–410. 10.1038/nature24268

34. Slaymaker, I. M., Gao, L., Zetsche, B., Scott, D. A., Yan, W. X., and Zhang, F. (2016). Rationally engineered Cas9 nucleases with improved specificity. Science 351, 84–88. 10.1126/science.aad5227

35. Kim, H. K., Lee, S., Kim, Y., Park, J., Min, S., Choi, J. W., Huang, T. P., Yoon, S., Liu, D. R., & Kim, H. H. (2020). High-throughput analysis of the activities of xCas9, SpCas9-NG and SpCas9 at matched and mismatched target sequences in human cells. Nat. Biomed. Eng. 4, 111–124. 10.1038/s41551-019-0505-1

36. Guo, L. Y., Bian, J., Davis, A. E., Liu, P., Kempton, H. R., Zhang, X., Chemparathy, A., Gu, B., Lin, X., Rane, D. A., Xu, X., Jamiolkowski, R. M., Hu, Y., Wang, S., and Qi, L. S. (2022). Multiplexed genome regulation in vivo with hyper-efficient Cas12a. Nat. Cell Bio. 24, 590–600. 10.1038/s41556-022-00870-7

37. Han, D., Xiao, Q., Wang, Y., Zhang, H., Dong, X., Li, G., Kong, X., Wang, S., Song, J., Zhang, W., Zhou, J., Bi, L., Yuan, Y., Shi, L., Zhong, N., Yang, H., and Zhou, Y. (2023). Development of miniature base editors using engineered IscB nickase. Nat. Methods 20, 1029–1036. 10.1038/s41592-023-01898-9

38. Kong, X., Zhang, H., Li, G., Wang, Z., Kong, X., Wang, L., Xue, M., Zhang, W., Wang, Y., Lin, J., Zhou, J., Shen, X., Wei, Y., Zhong, N., Bai, W., Yuan, Y., Shi, L., Zhou, Y., and Yang, H. (2023). Engineered CRISPR-OsCas12f1 and RhCas12f1 with robust activities and expanded target range for genome editing. Nat. Comm. 14, 2046. 10.1038/s41467-023-37829-7

39. Kato, K., Okazaki, S., Kannan, S., Esra Demircioglu, F., Isayama, Y., Ishikawa, J., Fukuda, M., Macrae, R. K., Nishizawa, T., Makarova, K. S., Koonin, E. V., Zhang, F., and Nishimasu, H. (2022). Structure of the IscB–ωRNA ribonucleoprotein complex, the likely ancestor of CRISPR-Cas9. Nat. Comm. 13, 6719. 10.1038/s41467-022-34378-3

40. Sasnauskas, G., Tamulaitiene, G., Druteika, G., Carabias, A., Silanskas, A., Kazlauskas, D., Venclovas, Č., Montoya, G., Karvelis, T., and Siksnys, V. (2023). TnpB structure reveals minimal functional core of Cas12 nuclease family. Nature 616, 384–389. 10.1038/s41586-023-05826-x

41. Saito, M., Xu, P., Faure, G., Maguire, S., Kannan, S., Vo, S., Desimone, A., Macrae, R. K., and Zhang, F. (2023). Fanzor is a eukaryotic programmable RNA-guided endonuclease. Nature 620, 660– 668. 10.1038/s41586-023-06356-2

42. Bepler, T., Morin, A., Rapp, M., Brasch, J., Shapiro, L., Noble, A.J., and Berger, B., (2019). Positive-unlabeled convolutional neural networks for particle picking in cryo-electron micrographs. Nat. Methods 16, 1153–1160. 10.1038/s41592-019-0575-8

43. Punjani, A., Zhang, H., and Fleet, D.J., (2020). Non-uniform refinement: adaptive regularization improves single-particle cryo-EM reconstruction. Nat. Methods 17, 1214–1221. 10.1038/s41592-020-00990-8

44. Emsley, P., and Cowtan, K., (2004). Coot: model-building tools for molecular graphics. Acta Crystallogr. D Biol. Crystallogr. 60, 2126–2132. 10.1107/S0907444904019158

45. Liebschner, D., Afonine, P.V., Baker, M.L., Bunkóczi, G., Chen, V.B., Croll, T.I., Hintze, B., Hung, L.W., Jain, S., McCoy, A.J., Moriarty, N.W., Oeffner, R.D., Poon, B.K., Prisant, M.G., Read, R.J., Richardson, J.S., Richardson, D.C., Sammito, M.D., Sobolev, O.V., Stockwell, D.H., Terwilliger, T.C., Urzhumtsev, A.G., Videau, L.L., Williams, C.J., and Adams, P.D., (2019). Macromolecular structure determination using X-rays, neutrons and electrons: recent developments in Phenix. Acta Crystallogr. D Struct. Biol. 75, 861–877. 10.1107/S2059798319011471

46. Afonine, P.V., Poon, B.K., Read, R.J., Sobolev, O.V., Terwilliger, T.C., Urzhumtsev, A., and Adams, P.D., (2018). Real-space refinement in PHENIX for cryo-EM and crystallography. Acta Crystallogr. D Struct. Biol. 74, 531–544. 10.1107/S2059798318006551

