## Supplemental Information for "Rapid DNA unwinding accelerates genome editing by engineered CRISPR-Cas9"

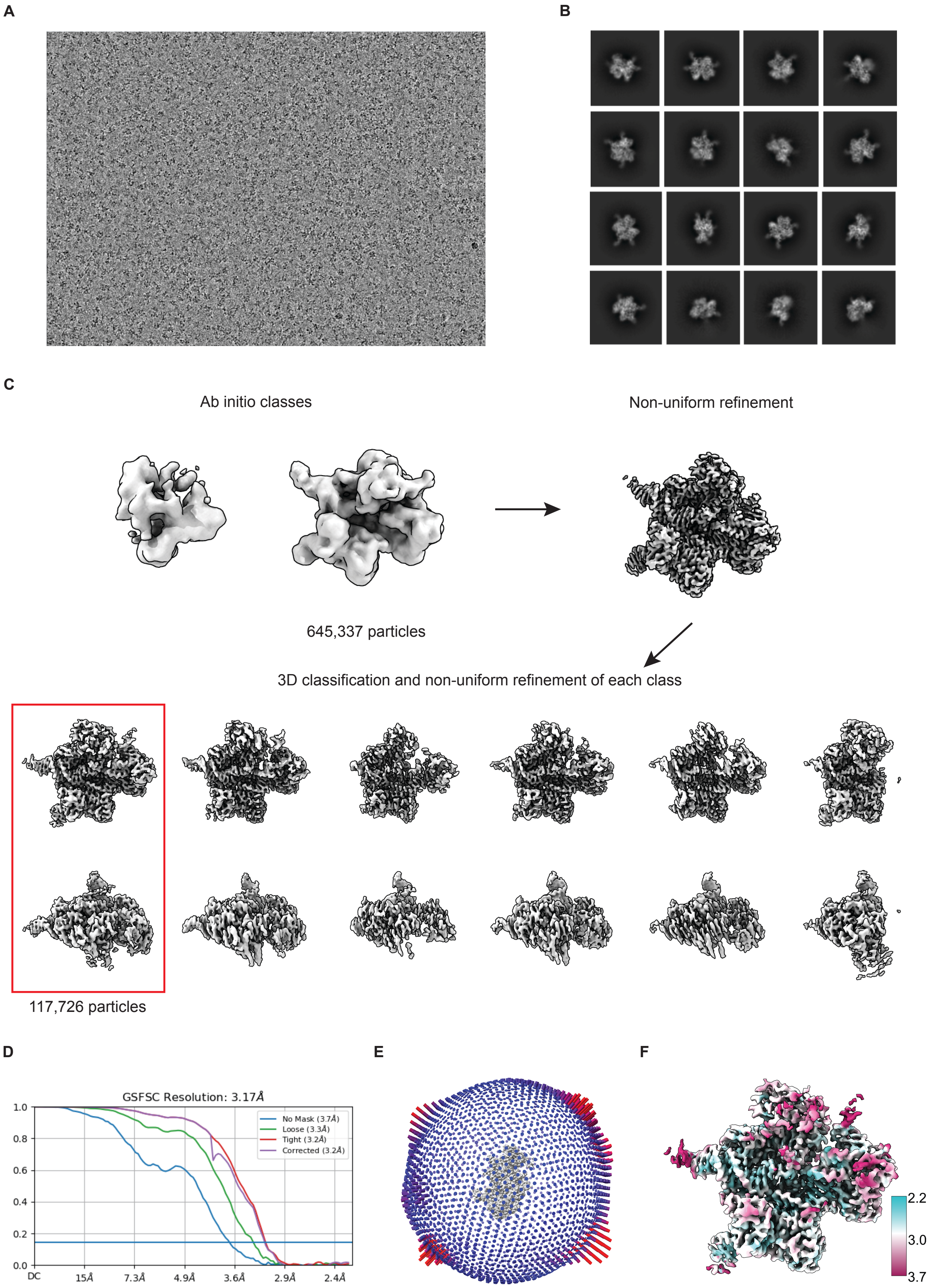
A
B
C
Ab initio classes
Non-uniform refinement
645,337 particles
3D classification and non-uniform refinement of each class
117,726 particles
D
GSFSC Resolution: 3.17Å
No Mask (3.7Å)
Loose (3.3Å)
Tight (3.2Å)
Corrected (3.2Å)
1.0
0.8
0.6
0.4
0.2
0.0
DC
15Å
7.3Å
4.9Å
3.6Å
2.9Å
2.4Å
E
F
2.2
3.0
3.7

**Figure S1:** Data processing schematic for wildtype GeoCas9, related to Figure 1

(A) Example cryo-EM image after beam-induced motion correction. (B) A subset of selected 2D class averages. (C) Cryo-EM data processing in cryoSPARC v.4.4. *Ab initio* classes are visualized at 0.2 contour level and refined classes at 0.27 contour level. Red box highlights the final map. (D) Gold standard FSC curves from the final round of non-uniform refinement in cryoSPARC. (E) Particle orientation distribution. (F) Local resolution map for final map calculated in cryoSPARC with threshold 0.143 and displayed in ChimeraX v. 1.6.1 with dust removal size 5 and contour level 0.27.

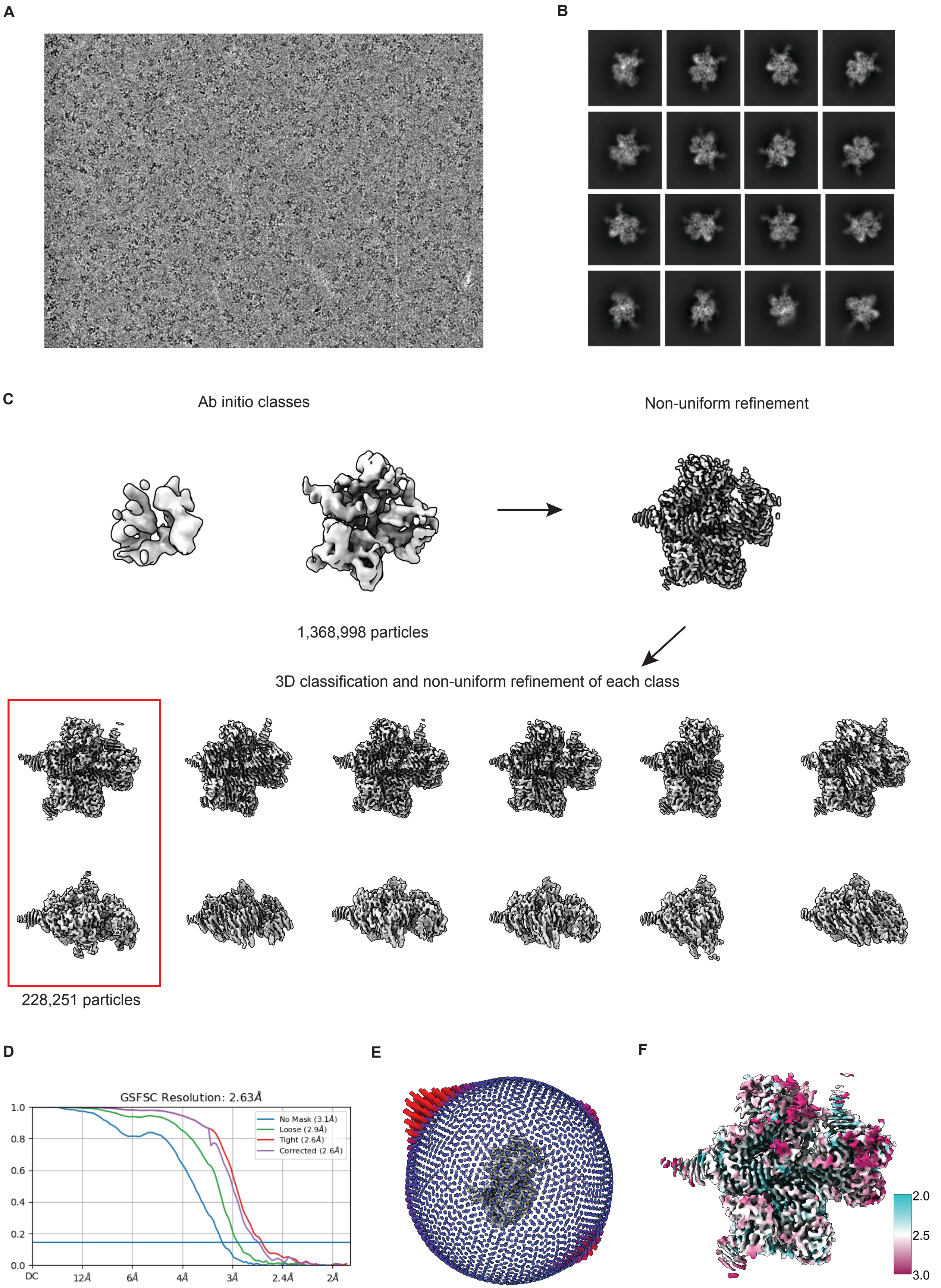
A
B
C
Ab initio classes
Non-uniform refinement
1,368,998 particles
3D classification and non-uniform refinement of each class
228,251 particles
D
GSFSC Resolution: 2.63Å
No Mask (3.1Å)
Loose (2.9Å)
Tight (2.6Å)
Corrected (2.6Å)
1.0
0.8
0.6
0.4
0.2
0.0
DC
12Å
6Å
4Å
3Å
2.4Å
2Å
E
F
2.0
2.5
3.0

**Figure S2:** Cryo-EM workflow for iGeoCas9, related to Figure 1

(A) Example cryo-EM image after beam-induced motion correction. (B) A subset of selected 2D class averages. (C) Cryo-EM data processing in cryoSPARC v.4.4. *Ab initio* classes are visualized at 0.2 contour level and refined classes at 0.27 contour level. Red box highlights the final map. (D) Gold standard FSC curves from the final round of non-uniform refinement in cryoSPARC. (E) Particle orientation distribution. (F) Local resolution map for final map calculated in cryoSPARC with threshold 0.143 and displayed in ChimeraX v. 1.6.1 with dust removal size 5 and contour level 0.27.

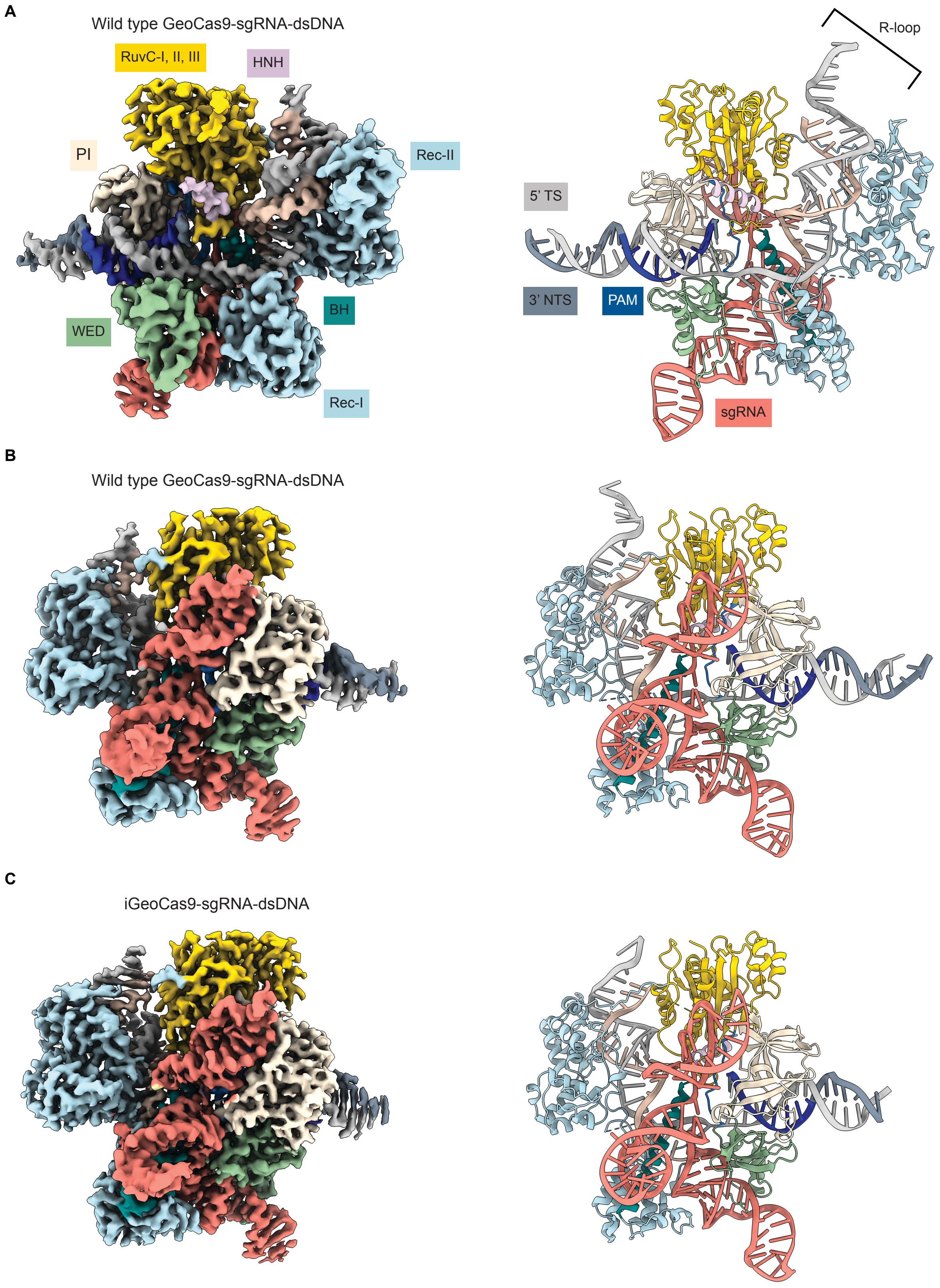
A
Wild type GeoCas9-sgRNA-dsDNA
RuvC-I, II, III
HNH
PI
Rec-II
WED
BH
Rec-I
R-loop
5' TS
3' NTS
PAM
sgRNA
B
Wild type GeoCas9-sgRNA-dsDNA
C
iGeoCas9-sgRNA-dsDNA

**Figure S3:** Cryo-EM density maps and models, related to Figure 1

**(A)** Cyo-EM density map of wildtype GeoCas9-sgRNA-dsDNA complex (left) at 3.17 Å resolution with domains labeled. Wildtype GeoCas9 Model complex (right) with nucleic acids labeled. **(B)** Cyo-EM density map and model of wildtype GeoCas9-sgRNA-dsDNA complex rotated 180° from part (A). **(C)** iGeoCas9sgRNA-dsDNA complex density map at 2.63 Å resolution and corresponding model. Both are rotated 180° from Fig. 1.

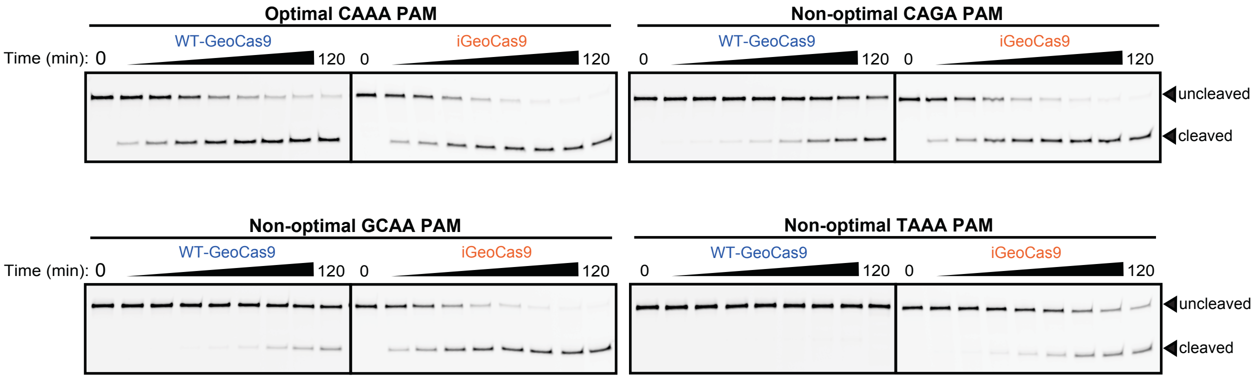


**Figure S4** iGeoCas9 exhibits broader PAM preferences *in vitro* than WTGeoCas9, related to Figure 3

In vitro dsDNA cleavage of wildtype (WT) GeoCas9 compared to iGeoCas9 using denaturing PAGE. 60 nucleotide substrates are 5’ 6-FAM labeled. PAM contained in each substrate listed above gel images. Fractions were collected at 0 sec, 30 sec, 1 min, 2.5 min, 5 min, 10 min, 30 min, 1 h, and 2 h.

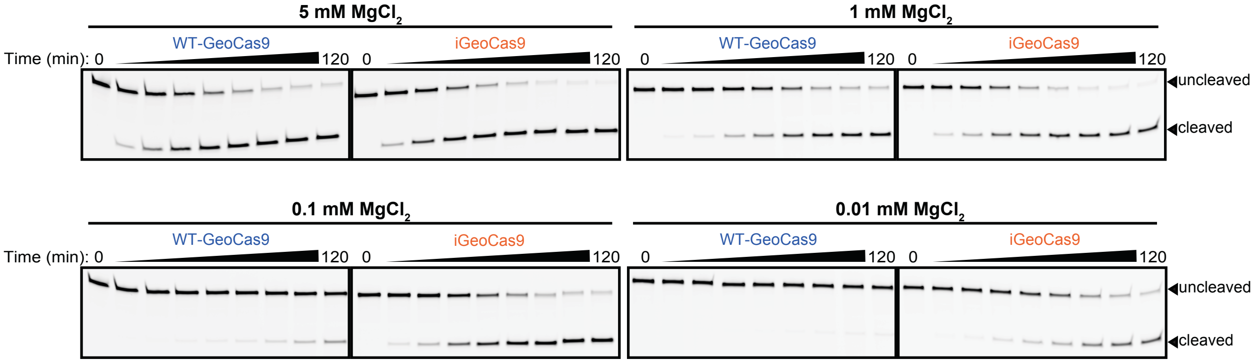


**Figure S5** The impact of $MgCl_2$ on WT-GeoCas9 and iGeoCas9 activity, related to Figure 5

In vitro dsDNA cleavage of wildtype (WT) GeoCas9 compared to iGeoCas9 using denaturing PAGE. 60 nucleotide substrates are 5'-6-FAM labeled and contain optimal PAM 5'-$N_4$CAAA-3'. $MgCl_2$ concentrations for each experiment indicated above gel images. Fractions were collected at 0 sec, 30 sec, 1 min, 2.5 min, 5 min, 10 min, 30 min, 1 h, and 2 h.

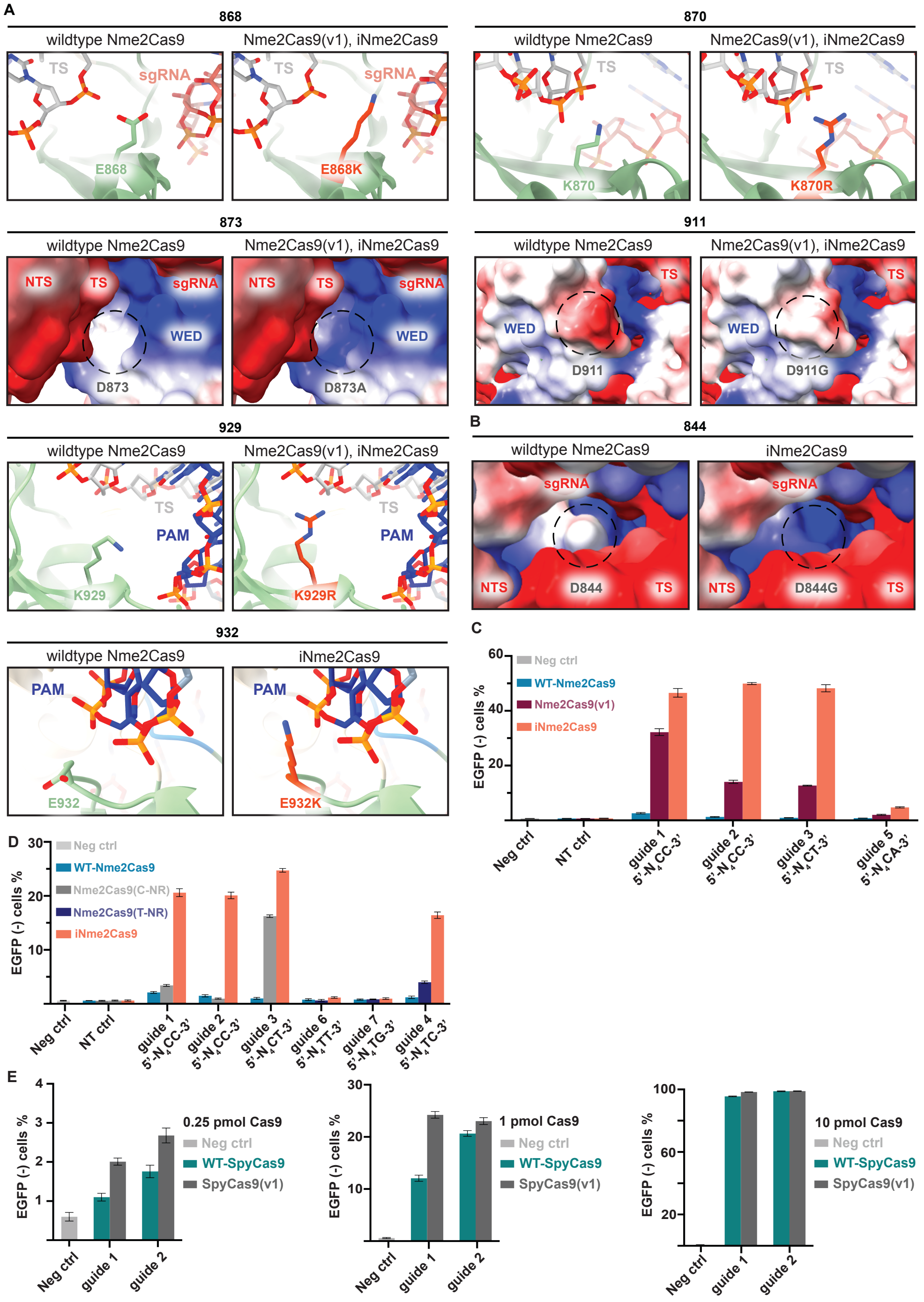


**Figure S6** Improved genome editors Nme2Cas9(v1) and iNme2Cas9 compared to wildtype Nme2Cas9, related to Figure 6, and preliminary engineering results of SpyCas9

**(A)** Structural comparison of wildtype Nme2Cas9 (PDB:6jE3)[1] and mutations contained in NmeCas9(v1) and iNmeCas9. Residue number located above images. Nme2Cas9(v1), iNme2Cas9 mutations were created in silico using ChimeraX (v1.6.1) using the wildtype Nme2Cas9 model and rotamers were chosen to demonstrate potential DNA interactions. Mutations expected to interact with DNA are represented as sticks and ribbons. Mutations expected to alter protein charge are represented as electrostatic potential maps (red, negative; blue, positive; white, non-polar). **(B)** Structural comparison of wildtype Nme2Cas9 (PDB:6jE3)[1] and mutations contained in iNmeCas2. **(C)** HEK293T cell editing by wildtype (WT-) Nme2Cas9, Nme2Cas9(v1), iNme2Cas9, with 6 different guides (PAM sequence below corresponding samples). Neg ctrl, no treatment control; NT, non-targeting guide control. **(D)** HEK293T cell editing by wildtype (WT-) Nme2Cas9, iNme2Cas9, Nme2Cas9(C-NR), and Nme2Cas9(T-NR) with 6 different guides (PAM sequence below corresponding samples). Neg ctrl, no treatment control; NT, non-targeting guide control. **(E)** HEK293T cell editing based on nucleofection of SpyCas9 RNPs (wildtype SpyCas9 or SpyCas9(v1) with 2 guides). Mutations in SpyCas9(v1) relative to WT-SpyCas9: G1104R, D1117A, D1125A, A1285K. Neg ctrl, no treatment control.

| Data collection | | |
|---|---|---|
| sample | Wildtype GeoCas9 | iGeoCas9 |
| Electron Microscope | FEI Talos Arctica | Titan Krios G2 |
| Electron Detector | K3 Direct Electron Detector | K3 Direct Electron Detector |
| Magnification | 36,000 | 81,000 |
| Voltage (kV) | 200 | 300 |
| Electron dose (e-/Å$^2$) | 50 | 50 |
| Defocus range (μm) | -0.8 to 2 | -0.8 to 2 |
| Pixel size (Å) | 0.57 | 0.465 |
| **3D reconstruction** | | |
| sample | Wildtype GeoCas9 | iGeoCas9 |
| Raw images | 2,767 | 7,849 |
| Initial particles | 1,026,723 | 6,498,580 |
| Final particles | 117,726 | 228,251 |
| Map resolution (Å) | 3.17 | 2.63 |
| FSC threshold | 0.143 | 0.143 |
| **Model refinement** | | |
| sample | Wildtype GeoCas9 | iGeoCas9 |
| Initial model used | Ab initio ModelAngelo model<br>Ab initio Colabfold model | Ab initio ModelAngelo model<br>Ab initio Colabfold model |
| Model resolution | 3.1 | 2.6 |
| FSC threshold | 0.143 | 0.143 |
| Model composition<br>Nonhydrogen atoms<br>Protein residues<br>Nucleotide<br>Ligands | <br>11072<br>910<br>171<br>0 | <br>10882<br>910<br>162<br>0 |
| B factors (mean, Å$^2$)<br>Protein<br>Nucleotide | <br>65.91<br>92.98 | <br>54.29<br>69.52 |
| R.m.s deviations<br>Bond length (Å)<br>Bond angles (°) | <br>0.003<br>0.509 | <br>0.003<br>0.528 |
| **Validation** | | |
| sample | Wildtype GeoCas9 | iGeoCas9 |
| MolProbity score | 1.71 | 1.55 |
| Clash score | 7.12 | 5.68 |
| Rotamer outlier (%) | 0 | 0 |
| Ramachandran statistics (%)-<br>Favored<br>Allowed<br>Outlier | <br>95.45<br>4.55<br>0 | <br>96.34<br>3.66<br>0 |
| Rama-Z score, whole<br>(r.m.s Rama-Z) | -0.07<br>0.28 | 0.50<br>0.28 |
| Map CC (box) | 0.76 | 0.67 |
| Map CC (mask) | 0.82 | 0.78 |

**Table S1** Cryo-EM data collection, 3D reconstruction, model refinement and validation, related to Figure

1

### SUPPLEMENTAL REFERENCES

1. Sun, W., Yang, J., Cheng, Z., Amrani, N., Liu, C., Wang, K., Ibraheim, R., Edraki, A., Huang, X., Wang, M., Wang, J., Liu, L., Sheng, G., Yang, Y., Lou, J., Sontheimer, E. J., and Wang, Y. (2019). Structures of *Neisseria meningitidis* Cas9 complexes in catalytically poised and anti-CRISPR-inhibited states. Mol. Cell *76*, 938–952. https://doi.org/10.1016/j.molcel.2019.09.025
